# stEDGE enables edge-guided multiscale reconstruction of hierarchical spatial domains and transition interfaces in spatial transcriptomics

**DOI:** 10.64898/2026.09.15.751915

**Authors:** Yi He, Shuqi Chen, Liqing Ding, Ruimin He, Xiaoqing Peng, Guihua Duan, Honglin Zhu, Hong-Dong Li, Shaokai Wang, Jianxin Wang

## Abstract

Spatial transcriptomic technologies enable high-resolution mapping of tissue architecture, yet most computational methods still represent tissues as flat spatial partitions, limiting their ability to resolve spatial domain boundaries, gradual transition regions and nested hierarchies. Here we present stEDGE, an edge-guided and interpretable framework for multiscale reconstruction of spatial domain hierarchies and transition-associated states in spatial transcriptomics. Unlike conventional domain-first approaches, stEDGE first estimates local boundary structure and uses it to guide the reconstruction of fine-grained spatial domains. We further introduce a domain transition index (DTI) to quantify domain-level transition propensity and integrate DTI with inter-domain similarity and boundary strength to organize fine domains into a coherent multilevel hierarchy. A tree-guided gene attribution strategy then distinguishes shared parent-level programs from branch-specific specialization across hierarchical spatial states. Across 15 benchmarked tissue sections spanning diverse tissues, technologies and spatial resolutions, stEDGE reconstructs developmental, inflammatory, tumor and brain architectures beyond conventional domain partitioning. Applied to cell-resolved Xenium profiling of fibrotic human lung, stEDGE further reveals lesion-associated remodeling as a coherent multiscale hierarchy in which remodeling interfaces coexist with a stable macrophage-associated airway/lumen state and spatially organized TLS-like lymphoid niches. Together, stEDGE provides a unified framework for modeling how stable compartments, domain boundaries, remodeling interfaces, localized niches and multiscale gene programs are jointly organized across spatial scales.

## Introduction

Spatial transcriptomic technologies have enabled transcriptome-wide profiling of intact tissues while preserving spatial context, with platforms ranging from array-based spatial transcriptomics to near-cellular and imaging-based assays.^1,2^ Accumulating ST studies have shown that tissue organization comprises discrete compartments, sharp domain boundaries, graded transition zones and nested structures across biological scales, which underlie diverse biological processes such as developmental patterning, inflammatory remodeling and complex tissue microenvironments. However, these structural features are not uniformly reflected in global transcriptomic differences. While some regions exhibit strong expression contrasts, others are separated by subtle or gradual changes, and boundary-localized signals may not correspond to globally separable clusters. As a result, accurately resolving spatial organization from ST data is inherently challenging, particularly in the presence of transition-associated regions and heterogeneous structural scales.

Most existing computational methods approach this problem through spatial domain identification. Traditional clustering frameworks developed for single-cell transcriptomic analysis (e.g., Seurat^3^, SCANPY^4^ and Louvain^5^) lack spatial awareness and often produce fragmented or biologically inconsistent assignments when applied to ST data. To address these limitations, a growing number of spatially aware methods have been developed to more accurately resolve tissue structure in space. Representative approaches (e.g., SpaGCN^6^, STAGATE^7^, GraphST^8^, SCAN-IT^9^, SEDR^10^, stLearn^11^ and STAIG^12^) are designed to recover locally coherent domains through graph-based learning, multimodal integration or spatially informed embeddings, whereas more recent methods (e.g., PROST^13^, STAMP^14^, SAGE^15^ and GASTON^16^) place greater emphasis on interpretable spatial features, latent programs or continuous topographic variation.

Despite these advances, three fundamental challenges persist in reconstructing spatial organization from ST data. First, explicitly representing spatial domain boundaries and biologically meaningful transition regions remains difficult. In many biological contexts, neighboring compartments may be separated by sharp domain boundaries or connected through finite-width transition zones enriched for intermediate or mixed states. However, many representative graph-based methods primarily rely on local aggregation over spatial proximity graphs, which favors locally coherent partitions while assigning each location to a single discrete domain. As a result, such discrete-label formulations may absorb finite-width transition zones into neighboring compartments or split them across hard boundaries, particularly when intermediate states are spatially broad or transcriptionally gradual. A more recent method, GASTON, models continuous spatial variation through an isodepth-based topographic formulation, but such a representation may be less flexible when multiple local transition axes coexist.

Second, tissue architecture is inherently multiscale, yet most existing methods operate at a single resolution or recover different granularities only through post hoc adjustment of clustering parameters. Although resolution tuning can generate partitions at different scales, it does not establish parent–child relationships or quantify transition strengths between spatial states. Broad tissue compartments often coexist with finer subdomains and localized microstates, requiring a representation that jointly captures structure across scales. Hierarchical clustering provides a natural framework for this purpose, but conventional hierarchies are typically constructed through sequential merging and therefore remain sensitive to the initial partition and early merge decisions. Recent single-cell methods, such as scSHC^17^ and CHOIR^18^, seek to improve hierarchical reliability through statistical testing and model-based pruning, but remain defined in transcriptomic space and do not incorporate the spatial constraints required for reconstructing tissue organization. Moreover, the resulting trees may not naturally align with interpretable biological levels such as compartments, domains and microenvironments.

Third, biological interpretation is commonly decoupled from spatial reconstruction and rarely aligned with multiscale spatial structure. Many deep learning–based methods, including graph neural networks and variational embedding frameworks^6–12^, rely on post hoc differential expression analysis to interpret latent representations, obscuring how individual genes contribute to spatial state organization. Recent interpretable approaches, such as STAMP and SAGE, associate spatial topics, domains or latent programs with informative gene modules, but do not directly organize these programs along a hierarchy of spatial states. Consequently, they cannot readily distinguish genes defining a shared parent-level identity from those specifying child-level branch specialization, limiting biological interpretation of how multiscale spatial states are organized and diversified. This distinction becomes particularly important when spatial states share a common macro-identity but differ in local activation, stromal interaction or microenvironmental context.

To address these challenges, we propose stEDGE, an edge-guided framework for reconstructing multiscale spatial organization in spatial transcriptomics. To our knowledge, stEDGE is the first unified framework in which local boundary structure directly guides fine-grained spatial domain reconstruction and is carried forward into subsequent multiscale hierarchical organization. Unlike conventional domain-first approaches that infer boundaries only after clustering, stEDGE first estimates local boundary signals and uses them to guide boundary-constrained expansion from stable low-boundary cores. We further introduce a domain transition index (DTI) that quantifies domain-level transition propensity and characterizes the continuum from stable compartments to transition-prone or structurally mixed spatial states. Together with inter-domain similarity and boundary strength, DTI guides the organization of fine domains into a coherent multilevel hierarchy. Finally, we introduce a tree-guided gene attribution strategy that links molecular programs to hierarchical spatial states, distinguishing shared parent-level identity from branch-specific specialization. Together, these components characterize spatial hierarchy, domain boundaries, transition-associated states and multiscale gene programs.

Through comprehensive benchmarking and applications across diverse tissues, technologies and spatial resolutions, stEDGE demonstrates that spatial organization cannot be fully captured by flat domain partitions alone. It resolves developmental boundaries, inflammatory transitions, tumor hierarchies and brain architectures through a shared boundary-aware multiscale representation. In cell-resolved Xenium profiling of fibrotic human lung, this framework further reveals lesion-associated remodeling as a coherent multiscale hierarchy, in which remodeling interfaces coexist with a stable macrophage-associated airway/lumen state and spatially organized TLS-like lymphoid niches. Together, stEDGE provides a unified framework for modeling how stable compartments, remodeling interfaces, localized niches and multiscale gene programs are jointly organized.

## Results

### Overview of stEDGE workflow

stEDGE is a computational framework for resolving fine-grained and multiscale tissue architecture from spatial transcriptomics data. Existing methods typically resolve spatial domains by clustering under a single global resolution, with boundaries treated as secondary by-products of partitioning. Yet anatomically meaningful domains are not always best defined by global transcriptomic separability, and some remain difficult to resolve under coarse clustering because their overall expression differences are modest. Inspired by image segmentation, in which stable interiors and explicit boundaries jointly define structure, stEDGE models spatial domain boundaries as primary structural signals rather than passive by-products of clustering. By leveraging boundary structure and transition relationships between adjacent regions, the framework resolves fine domains, organizes them into a spatial hierarchy and links this hierarchy to multiscale gene programs (Fig. 1a and Methods).

**Figure 1.**
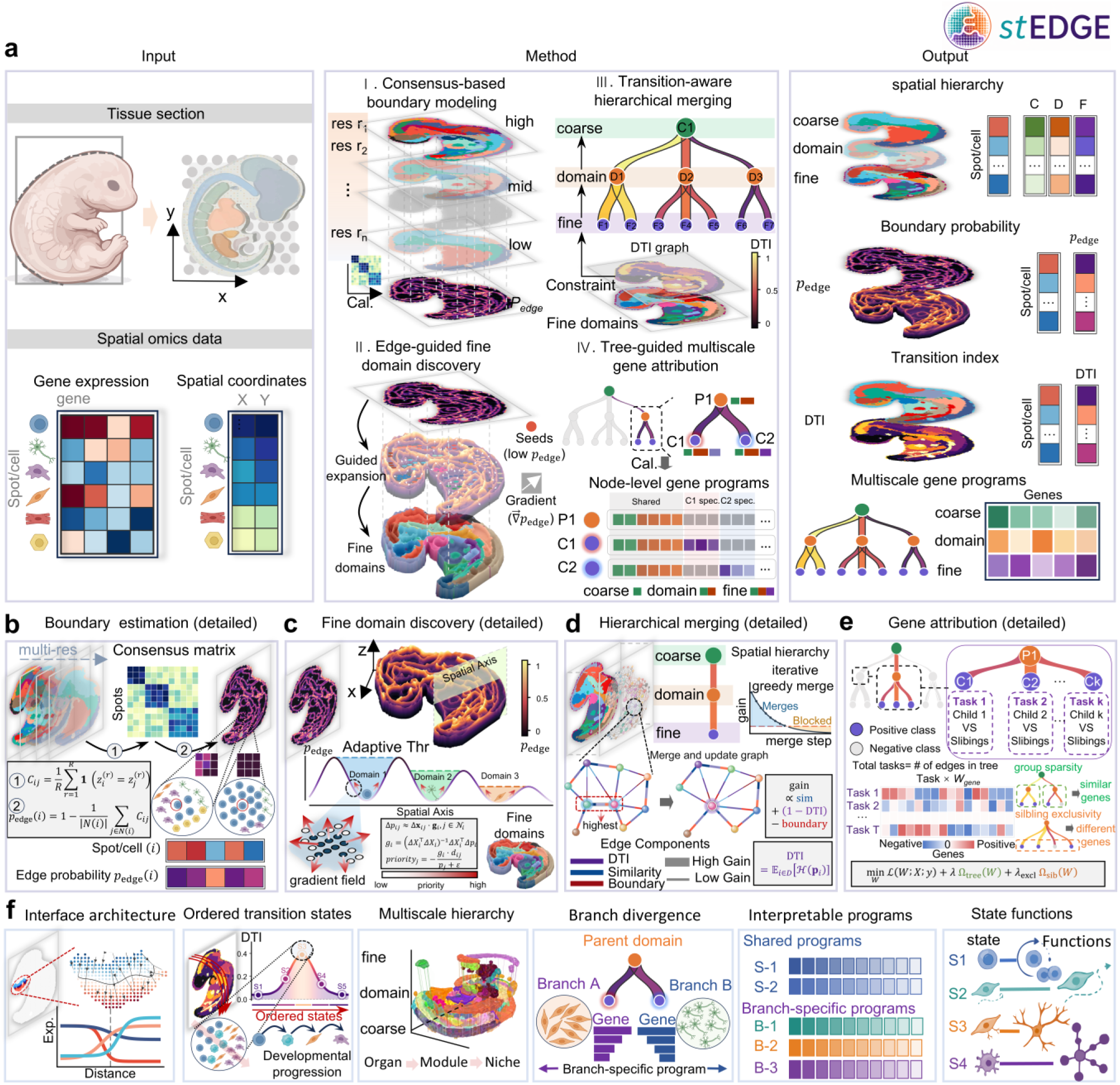
Overview of the stEDGE framework. a, stEDGE takes spatial transcriptomics data comprising gene expression and spatial coordinate matrices as inputs (left). The framework consists of four key modules (middle), including (I) consensus-based boundary modeling, (II) edge-guided fine-domain discovery, (III) transition-aware hierarchical merging and (IV) tree-guided multiscale gene attribution. These steps together produce structured outputs (right), including multiscale spatial hierarchy labels, boundary probability, transition index and multiscale gene programs. **b–e,** technical details of consensus-based boundary modeling (**b**), edge-guided fine-domain discovery (**c**), transition-aware hierarchical merging (**d**) and tree-guided multiscale gene attribution (**e**). **f,** Biological insights enabled by stEDGE, including interface architecture, ordered transition states, multiscale hierarchy, branch divergence, gene programs and state functions.

The stEDGE workflow comprises four integrated computational steps. First, stEDGE estimates a continuous boundary probability field using a consensus-based boundary modeling strategy. This step is designed to distinguish stable domain interiors from candidate boundary regions before segmentation, thereby establishing an explicit representation of spatial domain boundaries. Second, we introduce an edge-guided expansion strategy that propagates domain labels over the spatial graph under local boundary and gradient constraints. This step ensures that each domain is recovered according to the scale implied by its own boundary structure, rather than being constrained by a single global clustering resolution. We hypothesize that anatomically meaningful domains are often delineated by stable local boundary contrast, even when their overall transcriptomic separability is modest or heterogeneous. Such contrast may reflect differences in cellular composition and microenvironmental context across adjacent spatial domains. Third, to organize fine domains into higher-order tissue structure, stEDGE performs a transition-aware hierarchical merging procedure based on regional similarity, inter-domain boundary strength and domain-level transition propensity. This step ensures that the resulting coarse-to-fine hierarchy reflects structural relationships between neighboring domains, rather than being imposed by an arbitrary resolution parameter. Finally, stEDGE establishes multiscale gene programs for the learned spatial hierarchy through a tree-guided attribution framework based on parent–child relationships across hierarchical branches. This step ensures that the inferred programs simultaneously preserve shared transcriptional structure across related regions and identify branch-specific programs underlying divergence between neighboring spatial states.

stEDGE produces four classes of structured outputs, including multiscale spatial hierarchy labels, boundary probability, transition indices and multiscale gene programs. These outputs support downstream analyses of interface architecture, ordered transition states, multiscale hierarchy, branch divergence, gene programs and state functions (Fig. 1f). Selected domain boundaries may be further interpreted as biological interfaces or transcriptional transition zones when supported by anatomical context and continuous molecular variation. Together, stEDGE provides a unified framework for resolving fine domains, organizing tissue hierarchy and interpreting spatial architecture across biological scales.

### stEDGE resolves directional boundary transitions and cryptic neural crest states in the mouse embryo

Mouse embryo organogenesis provides a stringent benchmark for boundary-aware spatial reconstruction, as developing tissues comprise closely apposed anatomical compartments, lineage transitions and finite-width interfaces. We therefore evaluated stEDGE on nine MOSTA mouse embryo sections spanning E9.5 to E10.5, using manually curated anatomical annotations from Chen *et al.*^2^ (Cell, 2022) as reference labels (Fig. 2a; Supplementary Figs. 1–9). We compared stEDGE with 11 representative spatial domain or spatial structure inference methods: SAGE, SpaGCN, stLearn, GraphST, STAGATE, STAMP, PROST, SCAN-IT, STAIG, SEDR and GASTON. Across sections, stEDGE achieved the highest median clustering accuracy by ARI (0.342), NMI (0.587), HOM (0.623) and COM (0.560), and also showed the strongest boundary localization performance by AUPRC (0.686) and AUROC (0.737). Although GASTON yielded the highest median average silhouette width (ASW), stEDGE maintained competitive spatial continuity while more accurately matching anatomical labels and annotation-derived boundary labels (Supplementary Tables 3–10; Supplementary Data 1). In the representative E9.5_E1S1 section, stEDGE achieved the highest NMI (0.630) and ARI (0.287) and reconstructed fine-grained domains that closely followed manually annotated embryonic compartments (Fig. 2b). These results establish stEDGE as a robust framework for recovering anatomically coherent fine-grained domains and their boundaries across developing mouse embryo sections.

**Figure 2.**
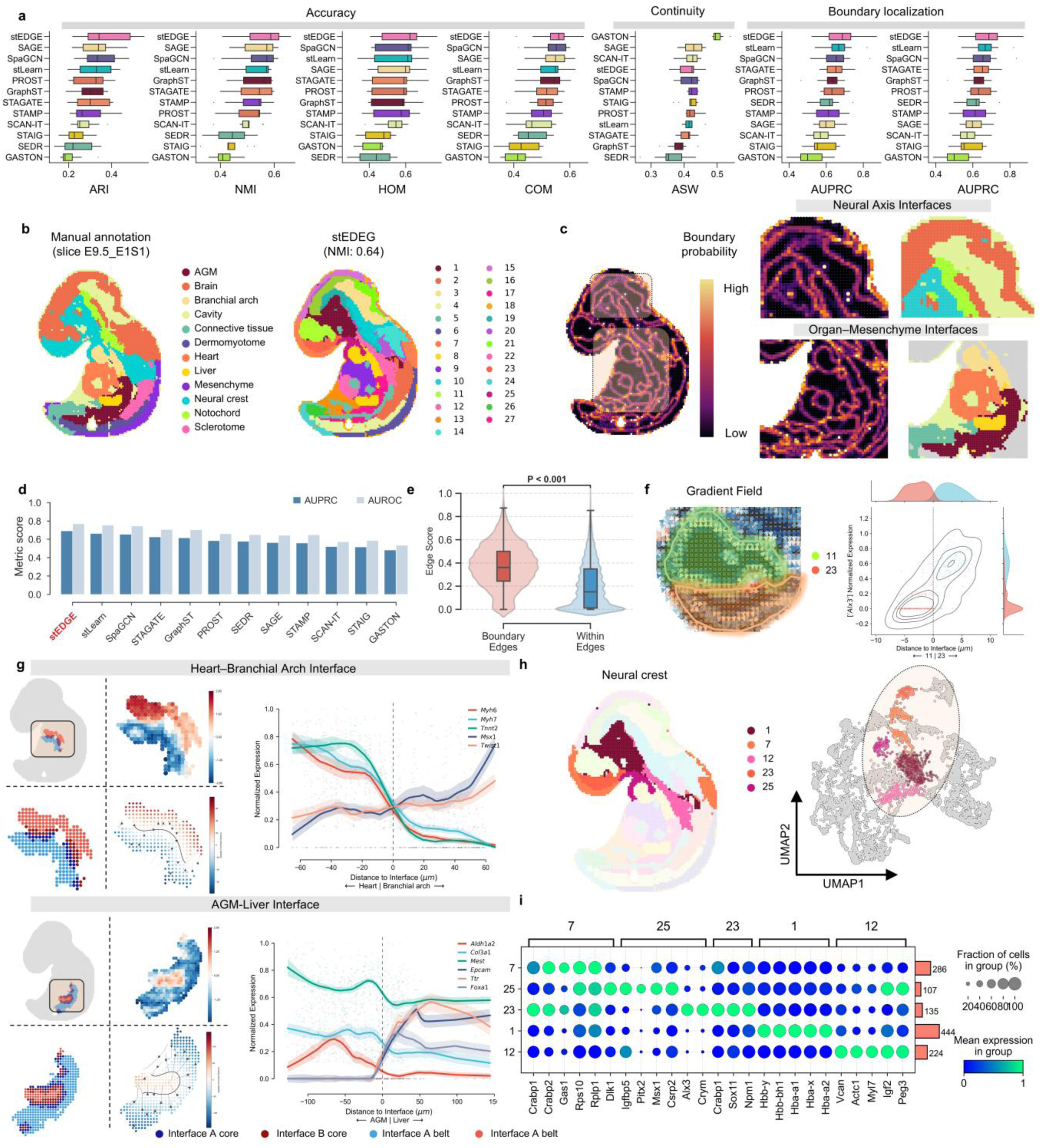
stEDGE reconstructs fine-grained spatial domains and localizes developmental boundaries in the mouse embryo. a, Benchmark performance of stEDGE and comparison methods across mouse embryo sections. Boxplots summarize clustering accuracy (ARI, NMI, HOM and COM), spatial continuity (ASW) and boundary localization performance (AUPRC and AUROC). **b,** Manual annotation and stEDGE-inferred fine-grained spatial domains for the representative section E9.5_E1S1. **c,** Boundary probability map inferred by stEDGE for the same section, with enlarged views of representative neural axis and organ–mesenchyme interfaces. **d,** Boundary localization performance on the representative section, shown by AUPRC and AUROC for each method. **e,** Distribution of stEDGE boundary scores for ground-truth boundary edges and within-region edges. **f,** Left, gradient field of the boundary probability across the section. Right, representative distance-resolved expression density across the Domain 11–23 interface. **g,** Two anatomically defined interfaces, Heart– Branchial arch (top) and AGM–Liver (bottom). For each interface, the inset indicates the spatial location, and the remaining panels show the interface belt, bilateral marker patterns and cross-interface expression profiles as a function of signed distance to the interface. **h,** Neural crest-related subdomains identified by stEDGE, shown in spatial coordinates (left) and UMAP space (right). **i,** Dot plot of representative genes across the five neural crest-related subdomains. Dot size indicates the fraction of cells in each group and color indicates mean expression.

A central component of stEDGE is the explicit modeling of spatial boundaries as a continuous probability field, rather than treating boundaries as implicit label transitions. On the representative section E9.5_E1S1, the inferred boundary probability map highlights major developmental separations and closely followed annotated anatomical interfaces, including the neural axis and organ– mesenchyme boundaries (Fig. 2c). To quantitatively evaluate boundary localization, we compared methods using boundary-focused metrics derived from ground-truth annotation transitions. stEDGE achieved the highest AUPRC (0.693) and AUROC (0.768) among all methods (Fig. 2 a,d). Consistently, edge scores assigned by stEDGE were significantly higher on ground-truth boundary edges than on within-region edges (Mann-Whitney *P* = 6.17 × 10⁻⁹, rank-biserial correlation = 0.428; Fig. 2e). Similar trends were observed across sections (Supplementary Figs. 1–9; Supplementary Data 2). These results establish boundary probability as a quantitative and interpretable representation of spatial interfaces.

We next asked whether the boundaries identified by stEDGE correspond to genuine transcriptional transition zones rather than discrete label discontinuities. The gradient field derived from the boundary probability showed structured directional changes across interfaces, consistent with ordered spatial transitions rather than isolated breaks in domain assignment (Fig. 2f). To examine this directly, we analyzed two anatomically defined interfaces: Heart–Branchial arch and AGM–Liver. In both cases, stEDGE captures narrow interface belts separating adjacent regions, within which marker genes exhibit reciprocal and continuous changes as a function of signed distance (Fig. 2g). For example, cardiac markers (*Myh6*, *Myh7*, *Tnnt2*) decreased across the Heart–Branchial arch interface, whereas branchial arch markers (*Msx1*, *Twist1*) increased in the opposite direction. Similar bilateral expression patterns were observed at additional anatomical (Dermomyotome) and domain-defined interfaces (Supplementary Figs. 16–17; Supplementary Tables 11–12; Supplementary Data 3 and 4). A representative stEDGE-derived interface, Domain 11–23, corresponding to the Neural crest–Brain boundary, further showed a spatially narrow but transcriptionally graded transition, with distinct gene expression programs restricted to opposite sides of the interface (Supplementary Fig. 18). Quantification of boundary transition breadth showed substantial gene-to-gene variation, ranging from a median breadth of 1.52 for *Cnn2* to 27.76 for *Crabp1*, with intermediate values for *Vim* (4.80), *Sox2* (9.90), *Id4* (23.77) and *Col1a2* (24.04) (Supplementary Fig. 18d). These observations indicate that the analyzed developmental interfaces are better represented as finite-width, directionally organized transcriptional transition zones than as purely discrete label discontinuities^19^.

Beyond recovering known anatomical compartments, stEDGE revealed spatially organized heterogeneity within the broadly annotated neural crest region that was not resolved by the reference annotation. Within the neural crest, stEDGE identified five transcriptionally distinct subdomains with partially contiguous spatial distributions and clear separation in UMAP space (Fig. 2h). Notably, this spatial heterogeneity is consistent with the well-established developmental properties of neural crest cells^20^, which form regionally distinct and transcriptionally diverse populations undergoing progressive fate restriction during embryogenesis^21,22^. Rather than forming a uniform compartment, neural crest cells are known to segregate into regionally distinct subpopulations with different differentiation potentials and to traverse continuous transcriptional trajectories associated with migration and fate commitment^20^.

Consistent with this view, stEDGE resolves subdomains corresponding to distinct developmental states. These subdomains exhibit distinct gene expression patterns (Fig. 2i; Supplementary Table 13) and are associated with different functional programs. For example, one subdomain shows enrichment for cardiopharyngeal or cardiac mesenchymal programs, marked by genes such as *Vcan*^23^ and *Myl7*^24^, whereas another displays a cranial neural crest-like state with elevated biosynthetic activity, characterized by genes including *Alx3*^25^ and *Crabp1*^26^. Additional subdomains are associated with signaling-active and patterning-related states, including enrichment for developmental regulators such as *Msx1*^27^ and *Dlk1*^28^ (Supplementary Figs. 19–23; Supplementary Data 5–7).

The spatial juxtaposition of these transcriptionally distinct but related states provides a mechanistic basis for the transition structures captured by stEDGE. These results highlight the ability of stEDGE to resolve spatially organized transcriptional states that are not captured by manual annotation, revealing hidden developmental heterogeneity within the neural crest. The corresponding hierarchy, domain transition structure and gene programs are further detailed in Supplementary Figs. 10–15 and Supplementary Figs. 24–28.

### stEDGE maps ordered spatial transition series and contractile stromal niches in Crohn’s disease

To test whether stEDGE can reconstruct organized tissue scaffolds alongside their embedded transition zones, we evaluated the framework on highly remodeled tissues. Stricturing Crohn’s disease provides a relevant test case because inflammatory remodeling, fibrosis and epithelial–stromal reorganization generate layered and transition-rich tissue architectures. We applied stEDGE to two intestinal Visium sections from Kong *et al.*^29^ (*Nat Genet*, 2025), using the original Region and Cluster annotations as reference labels (Fig. 3a–c; Supplementary Figs. 29–30). At the coarse region-level, stEDGE demonstrated the highest concordance with reference annotations across both sections, ranking first in key clustering metrics. This performance remained robust at the finer cluster-level, where stEDGE led in ARI, NMI, and COM while maintaining competitive HOM scores (Supplementary Tables 14–15). Visually, stEDGE faithfully recapitulated the banded intestinal wall organization, preserving the layered scaffold at fine resolution while simultaneously resolving localized cellular heterogeneity. In contrast, comparison methods either over-smoothed these boundaries or generated fragmented partitions (Fig. 3c; Supplementary Figs. 29–30). Together, these results demonstrate that stEDGE accurately reconstructs complex, layered disease architecture across scales.

**Figure 3.**
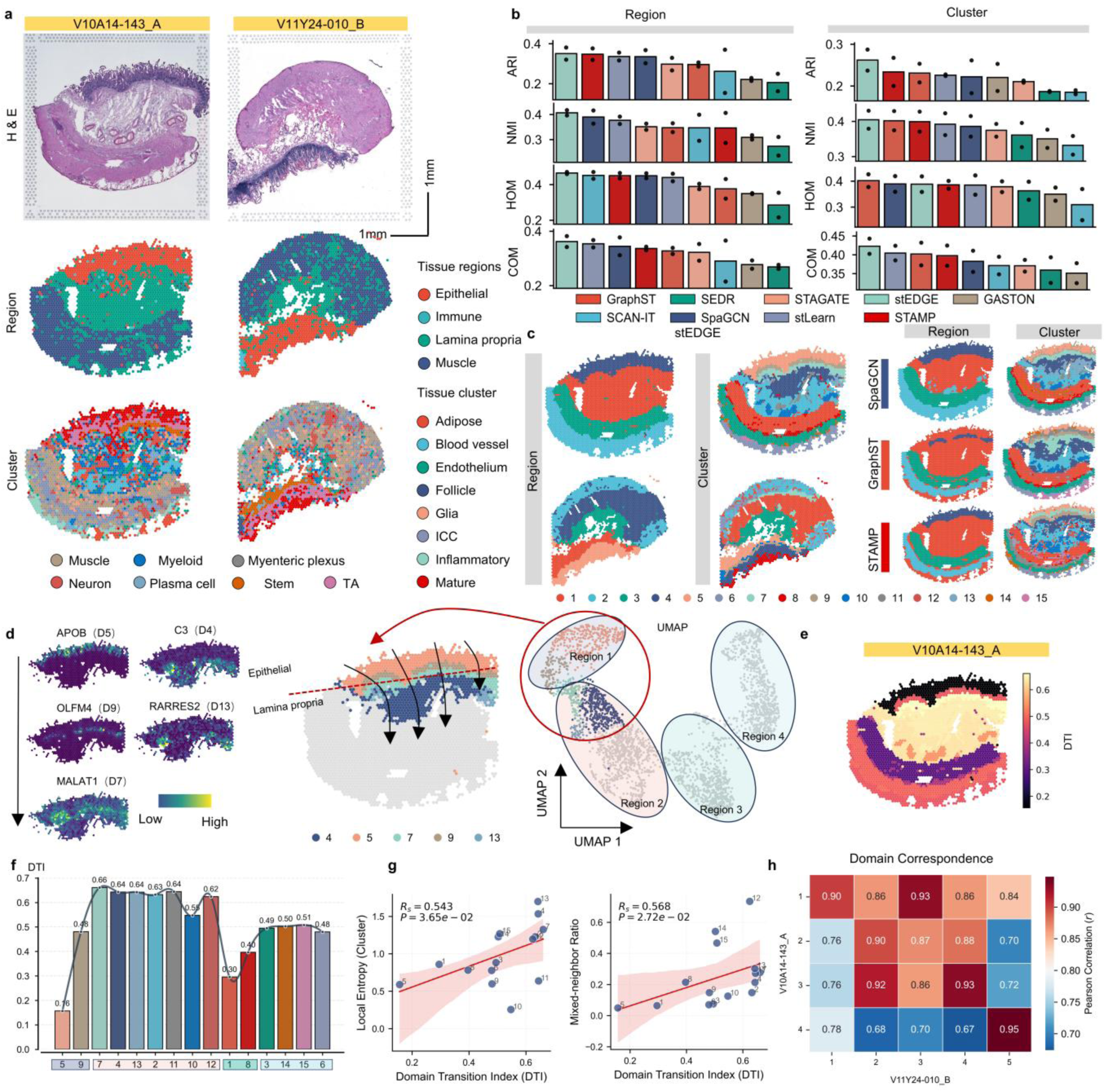
stEDGE maps ordered spatial transition series and contractile stromal niches in Crohn’s disease a, H&E images (top) and original region annotations (bottom) for sections V10A14-143_A and V11Y24-010_B. **b,** Benchmark performance of stEDGE and comparison methods across both sections. Boxplots summarize agreement with original labels at the coarse (Region) and fine (Cluster) levels using ARI, NMI, HOM, and COM (stEDGE is highlighted). **c,** Inferred spatial domain maps for both sections, with representative visual comparisons to other methods shown for V10A14-143_A. **d,** Characterization of the ordered transition domains (domains 4, 5, 7, 9, and 13) identified in V10A14-143_A. Panels show their representative marker genes (left), spatial locations (middle), and transcriptional relationships via UMAP projection (right). **e,** Spatial distribution of the DTI in the corresponding section. **f,** Mean DTI across domains ordered along the epithelial-to-deeper-tissue axis; boxes indicate the corresponding coarse domain partitions. **g,** Correlation of DTI with local entropy (left) and mixed-neighbor ratio (right). **h,** Cross-section correspondence of coarse domains between V10A14-143_A and V11Y24-010_B, illustrated by a Pearson-correlation heat map.

Having established reconstruction accuracy, we next examined whether stEDGE captures coherent spatial organization beyond discrete partitions. In the representative section V10A14-143_A, stEDGE resolved an ordered spatial transition series (5→9→7→4→13) that defines a continuous epithelial-to-stromal axis (Fig. 3d–f; Supplementary Figs. 31–34). This series progresses from a stable absorptive epithelial core (Domain 5), marked by *APOB*, *ALDOB*, *ANPEP* and *GUCA2A*, through a regenerative epithelial interface state (Domain 9), characterized by *OLFM4*, *REG1A*, *AGR2*, *PIGR* and *MUC2*, and a subepithelial mixed transition state (Domain 7), before reaching deeper stromal compartments enriched for fibroblast-associated genes (Domain 4 and Domain 13). Consistently, the domain transition index (DTI) increased along this series, rising from 0.16 in Domain 5 to 0.48 in Domain 9 and peaking at 0.66 in Domain 7, and remaining elevated in deeper stromal-like states (Fig. 3f). DTI further correlated positively with local entropy (Rₛ = 0.543, *P* = 3.65 × 10⁻²) and mixed-neighbor ratio (Rₛ = 0.568, *P* = 2.72 × 10⁻²) (Fig. 3g), indicating that higher DTI captures increased local heterogeneity and state mixing. These results show that stEDGE organizes Crohn’s disease tissue into stable cores and transition-associated states along a coherent spatial axis.

We next asked whether these transitions correspond to continuous molecular changes rather than discrete domain shifts. Analysis of representative transitions along the series, including epithelial-to-interface (5|9), interface-to-subepithelium (9|7) and stroma-associated transitions (7|4), revealed coordinated and gradual changes in gene expression across spatial boundaries (Supplementary Fig. 34g; Supplementary Table 16; Supplementary Data 8 and 9). For example, mature epithelial genes such as *GUCA2A* and *MT-CO1* decreased across the 5|9 transition, whereas regenerative markers including *OLFM4* and *REG1A* increased toward the interface state. The 9|7 transition showed loss of secretory epithelial genes (*AGR2*, *PIGR*, *MUC2*) together with increased expression of mitochondrial genes (*MT-CO3*, *MT-CYB*), consistent with a metabolically active transition zone. The 7|4 transition exhibited a clear cross-compartment shift from epithelial-associated markers (*PIGR*, *REG1A*, *DEFA6*) to stromal genes such as *SFRP2* and *FTH1*. Across all transitions, marker genes displayed spatially concordant enrichment, smooth distance-resolved changes and coherent expression structure. These observations indicate that domain boundaries correspond to structured transcriptional gradients rather than discrete discontinuities.

Finally, we asked whether transition-aware metrics derived from stEDGE could reveal hidden heterogeneity within the lamina propria. Along the epithelial-to-stromal transition axis, domains with elevated DTI were not restricted to boundary regions but extended into deeper stromal compartments (Fig. 3f), suggesting the presence of transition-associated states beyond canonical interfaces^30^. Notably, a high DTI domain (Domain 13) was consistently localized adjacent to the subepithelial transition zone and occupied a spatially restricted region within the lamina propria (Supplementary Fig. 35a). Differential expression and functional enrichment suggest that this region represents a contractile stromal-like niche, characterized by a mesenchymal and myofibroblast-associated program^31^ including *TAGLN*, *MYL9*, *ACTG2*, *TPM1* and *TPM2*, together with stromal niche-associated genes such as *AEBP1*^32^, *RARRES2*, *COL6A* and *GREM1* (Supplementary Fig. 35b–d; Supplementary Data 10). Functional enrichment further supported this interpretation, highlighting pathways related to muscle contraction, cell–matrix adhesion and elastic fiber assembly^33^ (Supplementary Fig. 35e; Supplementary Data 11). In contrast, the neighboring lower DTI region corresponds to a broader fibroblast-rich stromal state^34^. In this dataset, high-DTI domains coincided with specialized stromal programs associated with transition-rich microenvironments, rather than representing homogeneous terminal compartments. Cross-section comparison further showed that this transition-associated organization is reproducible at the coarse-level despite variation in fine-scale composition (Fig. 3h; Supplementary Fig. 36). Together, these findings demonstrate that stEDGE not only reconstructs spatial architecture, but also enables the identification of niche-level heterogeneity through transition-aware structural metrics.

### stEDGE decodes hierarchical spatial states and branch-specific gene programs in human breast cancer

To assess whether stEDGE can reconstruct pathology-related spatial architecture within highly heterogeneous clinical tissues^35^, we benchmarked it against nine representative methods on a human breast cancer section using the pathology reference annotation as ground-truth (Fig. 4a–c; Supplementary Figs. 37–39; Supplementary Table 17). stEDGE showed the strongest overall agreement with the reference labels, ranking first in ARI (0.645), NMI (0.715), HOM (0.730) and COM (0.690). Visually, stEDGE reconstructed coherent coarse- and domain-level maps that follow major pathology-defined contours while preserving local structure around central interfaces; notably, the IDC_2-associated region was recovered as a continuous compartment. By contrast, PROST produced smoother but coarser partitions that blurred local boundaries, whereas GraphST yielded fragmented assignments with reduced continuity of large pathology-associated regions. Importantly, the inferred organization is not a simple restatement of the original pathology labels. Across fine, domain and coarse levels, stEDGE revealed a higher-order spatial scaffold with coherent marker gene patterns (Supplementary Fig. 38). The inferred coarse domains show strong but incomplete correspondence to both four-class and fine-grained pathology annotations, with IDC distributed across multiple coarse domains and tumor-edge signals spanning nearly all compartments (Supplementary Fig. 39). This organization is further supported by purity and entropy: coarse domains 2 and 4 exhibit high purity and low entropy, consistent with stable compartments, whereas coarse domain 1 shows low purity and high entropy, indicating a strongly mixed state. Together, these results demonstrate that stEDGE reconstructs pathology-aligned organization while revealing a higher-order spatial scaffold beyond annotation-defined regions.

**Figure 4.**
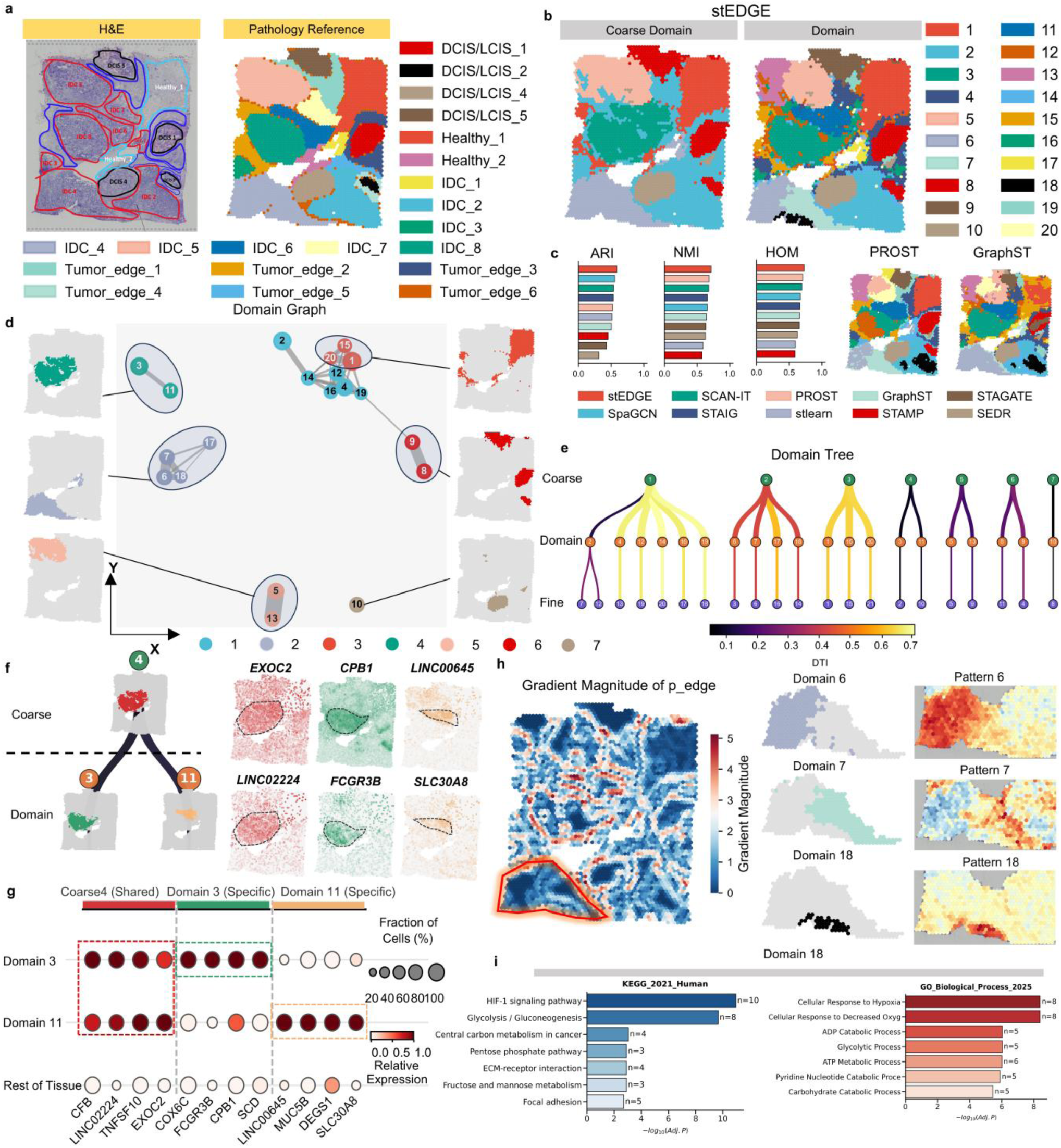
stEDGE decodes hierarchical spatial states and branch-specific gene programs in human breast cancer. a, H&E image and pathology reference annotation of the human breast cancer section. **b,** stEDGE-inferred spatial maps at the coarse domain and domain levels. **c,** Benchmark comparison of stEDGE and competing methods. Left, ARI, NMI and HOM scores. Right, representative spatial maps produced by PROST and GraphST. **d,** Domain graph of the stEDGE-inferred domains. Nodes represent domains, edges indicate adjacency relationships, and node colors indicate coarse domain membership. Side panels show the spatial extent of representative coarse compartments. **e,** Domain tree across the coarse, domain and fine levels. Tree-edge color indicates the DTI of the corresponding child domain. **f,** Tree-guided gene attribution for one branch of the hierarchy, centered on coarse 4 and its child domains 3 and 11. Representative top-weighted genes illustrate shared parent-level and branch-specific child-level programs. **g,** Bubble plot of representative genes for coarse 4, domain 3 and domain 11. Dot size indicates the fraction of cells or spots in each group and color indicates relative expression. **h,** Local transition-rich example from the lower-left region of the tissue. Left, gradient magnitude of *p*_edge_. Middle, spatial maps of domains 6, 7 and 18 within the same coarse compartment. Right, corresponding gene-pattern maps derived from the gene tree. **i,** Functional enrichment analysis of domain 18.

Tumor organization extends beyond pathology-defined regions and exhibits structured higher-order relationships (Fig. 4d,e; Supplementary Figs. 40–42). Domains form compact adjacency modules that are further organized into a non-uniform hierarchy. The domain graph shows that domains assigned to the same coarse compartment form compact adjacency modules rather than arbitrary collections of clusters. These modules include both simple domain pairs (for example, 3–11, 5–13 and 8–9) and larger assemblies such as the 6–7–17–18 module (Fig. 4d). Consistently, the domain tree organizes the tissue into a multiscale hierarchy spanning coarse, domain and fine states, with branches connected by tree edges associated with different DTI values (Fig. 4e). This hierarchy is strongly non-uniform across branches: some coarse domains show extensive internal branching (for example, coarse domain 1), whereas others form compact or near-linear structures (for example, coarse domain 7), with intermediate complexity observed in other branches (Supplementary Fig. 41). Branch strengths are likewise heterogeneous, with higher DTI edges concentrated in more elaborately subdivided regions and lower DTI edges marking compact structures. The stEDGE-inferred tumor organization therefore exhibits a structured and non-uniform hierarchical topology rather than a balanced or purely similarity-driven clustering tree.

This spatial hierarchy is accompanied by coordinated multiscale molecular programs across tumor branches (Fig. 4f,g; Supplementary Figs. 43–45). Tree-guided gene attribution assigns genes to specific hierarchical levels, enabling the separation of shared regional programs from branch-specific specialization. Focusing on a representative branch centered on coarse domain 4, we find that parent-level genes capture a shared regional program present across child domains 3 and 11, including *CFB*, *LINC02224*^36^, *TNFSF10*^37^ and *EXOC2*, consistent with a common inflammatory and signaling context. In contrast, child-level genes distinguish distinct local states within the same pathology-aligned region.

Domain 3 preferentially expresses *COX6C*, *FCGR3B*^38^, *CPB1*^39^ and *SCD*^40^, suggesting a local state with metabolic/lipid-associated and immune-adjacent features; notably, *CPB1* has been reported in ductal breast lesions^39^, although its functional relevance in this spatial context remains unclear. In contrast, Domain 11 expresses *LINC00645*, *MUC5B*^41^, *DEGS1* and *SLC30A8*, suggesting a localized secretory-like program. Spatial maps, dot plot summaries and expression comparisons confirm that parent-level genes mark shared compartments, whereas child-level genes highlight localized branch-specific programs (Fig. 4f,g; Supplementary Fig. 45). This hierarchical assignment is not directly provided by standard cluster-wise differential expression analysis, which typically treats clusters independently and does not explicitly model nested parent–child relationships.

To assess cross-section reproducibility, we projected the same 12-gene set defining the parent-shared and branch-specific programs onto a second breast cancer section without gene re-selection. Section 2 domain 4 (A-like) and domain 7 (B-like) showed the strongest correspondence to the Section 1 domain 3 and domain 11 states, respectively, and preserved the parent-shared program and reciprocal branch-specific enrichment (Supplementary Fig. 46; Supplementary Table 18). Gene-level effect estimates showed strong concordance across sections (Pearson’s (r = 0.962), (*P* = 5.66 × 10^−7^); Spearman’s (*ρ* = 0.923), (*P* = 1.86 × 10^−5^); 91.7% sign concordance), supporting the cross-section reproducibility of these tree-guided molecular programs.

Finally, we asked whether transition-aware structure reveals hidden local states within tumor compartments (Fig. 4h,i; Supplementary Figs. 47–53; Supplementary Data 12 and 13). Within a transition-rich compartment (coarse domain 2), stEDGE resolves multiple spatially adjacent but transcriptionally distinct states (domains 6, 7 and 18), each defined by branch-specific gene programs. Among these, domain 18 shows the clearest functional specialization, with elevated expression of *PGK1*, *GAPDH*, *ENO1* and *S100A6* and enrichment of hypoxia- and glycolysis-related pathways^42^, consistent with a localized glycolytic and stress-associated program^43,44^. Domains 6 and 7 represent additional sibling states within the same compartment, each with distinct discriminative gene patterns. Tree-guided classification confirms that these states are robustly separable (AUC = 0.964, 0.991 and 0.994; AP = 0.963, 0.986 and 0.954; Supplementary Figs. 50–52). More broadly, transition-prone regions are distributed across the tissue rather than confined to a single branch. Gradient-based hotspots form a spatial network, and DTI correlates positively with both local entropy (Spearman R = 0.63, P < 0.001) and mixed-neighbor ratio (Spearman R = 0.61, P < 0.001) (Supplementary Fig. 53), indicating that transition-associated structure is distributed across this tumor section rather than confined to a single branch^45^. Together, these results show that stEDGE integrates spatial hierarchy, transition structure and multiscale gene programs into a unified representation of tumor architecture, revealing hidden states beyond conventional pathology annotation.

### stEDGE achieves structure-agnostic generalization across region-based and layer-based brain architectures

Brain tissues provide a stringent test of whether spatial reconstruction is tied to a particular architectural regime. The mouse hippocampus is organized into regionally segregated subfields, whereas the cerebellum is defined by thin, folded and repetitive laminar structures. These contrasting geometries require a method to preserve local boundaries and marker-supported domains without assuming a single global tissue layout. We therefore evaluated stEDGE on high-resolution Slide-seqV2 hippocampus and cerebellum datasets to test whether it generalizes across region-based and layer-based brain architectures (Fig. 5a).

**Figure 5.**
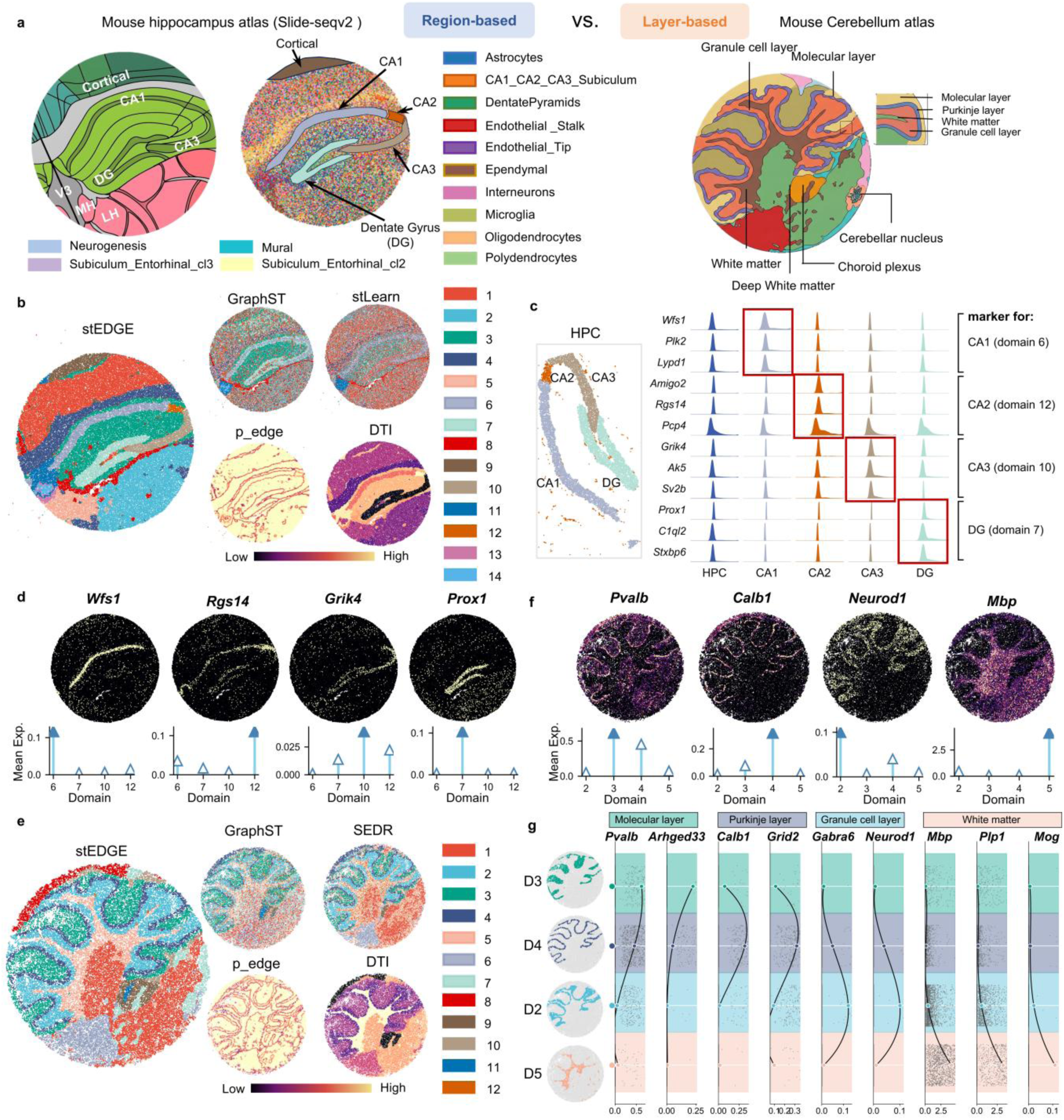
stEDGE achieves structure-agnostic generalization across region-based and layer-based brain architectures. a, Reference atlases and schematic spatial organization of the mouse hippocampus and cerebellum, representing region-based and layer-based architectures, respectively. **b,** Spatial domains inferred by stEDGE in the mouse hippocampus, with representative comparisons to GraphST and stLearn, together with p_edge and DTI maps. **c,** UMAP projection of hippocampal spots and ridge plots of representative marker genes across CA1-, CA2-, CA3- and dentate gyrus (DG)-associated domains. **d,** Spatial expression maps and mean expression summaries of representative hippocampal markers, including *Wfs1* (CA1), *Rgs14* (CA2), *Grik4* (CA3) and *Prox1* (DG). **e,** Spatial domains inferred by stEDGE in the mouse cerebellum, with representative comparisons to GraphST and SEDR, together with p_edge and DTI maps highlighting recurrent laminar interfaces. **f,** Spatial expression maps and mean expression summaries of representative cerebellar markers, including *Pvalb* (molecular layer), *Calb1* (Purkinje layer), *Neurod1* (granule cell layer) and *Mbp* (white matter). **g,** Layer-specific expression summaries across cerebellar domains D3, D4, D2 and D5, corresponding to the molecular layer, Purkinje layer, granule cell layer and white matter, respectively. Representative markers (*Pvalb, Arhged33*; *Calb1, Grid2*; *Gabra6, Neurod1*; *Mbp, Plp1, Mog*) support the inferred layer identities.

In the mouse hippocampus, a canonical region-based system with well-defined anatomical subfields, stEDGE accurately recovered spatial organization consistent with known neuroanatomy (Fig. 5b–d; Supplementary Figs. 54–58; Supplementary Tables 19–21). Across methods including GraphST, stLearn, SEDR and STAMP, stEDGE achieved the highest Silhouette score (0.171) and Spatial Coherence (0.936), together with the lowest Davies-Bouldin index (2.102) and the best Spatial Silhouette value (−0.113). The inferred domains follow the characteristic curved geometry of hippocampal subfields, preserving the continuity of CA regions and the dentate gyrus. Domain assignments correspond to canonical anatomical regions, with domains 6, 12, 10 and 7 mapping to CA1, CA2, CA3 and dentate gyrus, respectively, and supported by established marker genes, including *Wfs1*, *Rgs14*, *Grik4* and *Prox1*^46,47^. Region-specific transcriptional programs further reinforce these assignments, including calcium signaling and glutamatergic synapse-related functions in CA1-associated domains and synaptic organization in dentate gyrus-associated domains (Supplementary Figs. 59–66; Supplementary Data 14 and 15). These results demonstrate that stEDGE faithfully reconstructs regionally segregated spatial organization in a canonical brain system.

A more stringent test arises in the cerebellum, where spatial organization is defined by thin, highly folded and repetitive laminar structures (Fig. 5e–g; Supplementary Figs. 67–79; Supplementary Tables 20–21). Despite this complexity, stEDGE recovered continuous layered domains that remain coherent across repeated folds and preserve the arborized white matter scaffold. In contrast, GraphST exhibits increased mixing across adjacent laminae, stLearn fails to preserve the folded laminar pattern and SEDR produces broader but less sharply defined partitions. Quantitatively, stEDGE again achieved the strongest overall performance, with the highest Silhouette score (0.127) and Spatial Coherence (0.891), the lowest Davies-Bouldin index (2.183) and the second-best Spatial Silhouette value among compared methods. The inferred domains align with canonical cerebellar layers, including the molecular layer (domain 3), Purkinje layer (domain 4), granule cell layer (domain 2) and white matter (domain 5), supported by known markers such as *Pvalb* and *Arhged33*, *Calb1* and *Grid2*, *Gabra6* and *Neurod1*, and *Mbp*, *Plp1* and *Mog*^48,49^. Functional enrichment further supports these identities, with myelination-related programs enriched in white matter and synaptic transmission pathways in the granule cell layer^50^. This performance is consistent with the ability of stEDGE to model local boundary and transition structure independent of large-scale geometry.

Across these two systems, stEDGE recovers spatial organization through shared structural principles rather than dataset-specific tuning. In both hippocampal and cerebellar datasets, boundary probability (p_edge) and DTI trace anatomically meaningful interfaces, capturing curved regional boundaries in the hippocampus and recurrent laminar interfaces in the cerebellum (Supplementary Figs. 55–56 and Supplementary Figs. 68–69). These signals highlight transitions as structured spatial features rather than isolated discontinuities. Notably, the two systems represent complementary spatial regimes—region-based segregation and layer-based repetition—yet are reconstructed within the same analytical framework. Together, These two contrasting datasets support the ability of stEDGE to operate across distinct spatial regimes without tissue-specific architectural assumptions^51,52^.

### stEDGE resolves multiscale fibrotic lung remodeling from cell-resolved Xenium data

To evaluate whether stEDGE generalizes to cell-resolved, imaging-based spatial transcriptomics, we analyzed SSc_1_1_2, a fibrotic human lung section from a Xenium dataset of systemic sclerosis-associated interstitial lung disease (SSc-ILD) generated by Markov *et al.*^53^ (82,224 cells and 378 genes; Fig. 6a; Supplementary Fig. 80a). Fibrotic lungs present an exceptionally challenging test case for spatial reconstruction, as preserved alveoli, expanded interstitium, and immune-enriched niches are densely interwoven within a highly remodeled multicompartment architecture^54^. To establish a rigorous benchmark reference, we generated manual annotations at two complementary scales. At the structural level, we integrated multiplexed immunofluorescence imaging with tissue morphology to partition the section into nine reference regions spanning airway, alveolar, vascular, and pleural compartments. At the pathological level, we manually delineated the section into a non-lesion-like area (NLA) and a lesion-associated fibrotic remodeling area (LA) (Fig. 6b; Supplementary Fig. 80b; Methods). These expert-curated reference annotations provide a structural benchmark to evaluate stEDGE’s capacity to resolve cell-level, multiscale tissue states.

**Figure 6.**
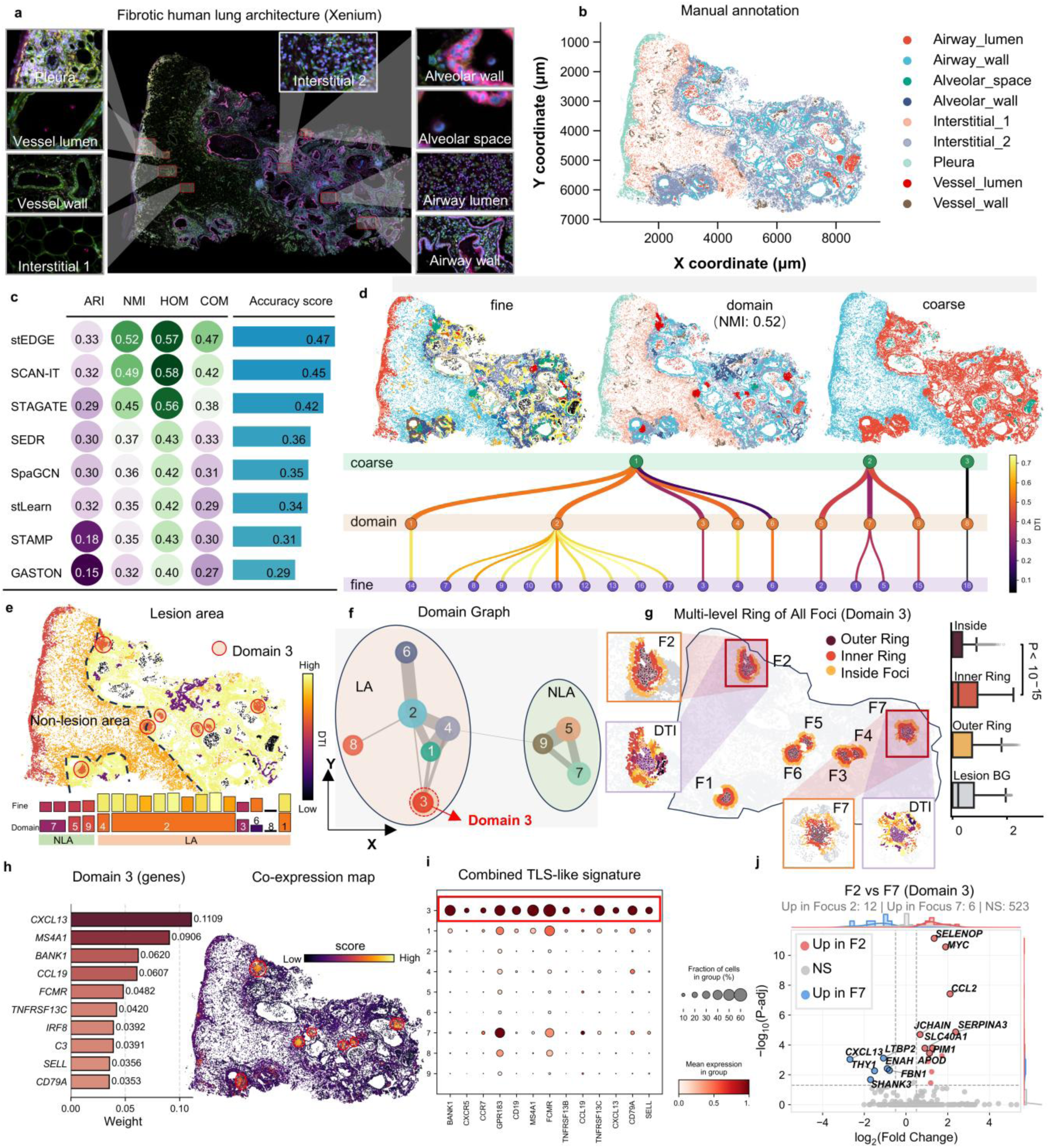
stEDGE resolves multiscale fibrotic lung remodeling from cell-resolved Xenium data, Xenium immunofluorescence image of the fibrotic human lung section, with enlarged representative views highlighting distinct anatomical and pathological structures. **b,** Manual reference annotation of the same section. **c,** Benchmark performance of stEDGE and comparison methods against the manual annotation. Bar plots summarize the clustering agreement using ARI, NMI, HOM, COM, and overall accuracy score. **d,** Hierarchical spatial organization inferred by stEDGE. Top, spatial maps at the fine, domain and coarse levels. Bottom, the domain tree linking the three hierarchical levels, with each tree edge colored according to the DTI of its child node. **e,** Spatial map of DTI across the tissue, distinguishing the lesion area (LA) from the NLA. Red circles highlight the stable Domain 3 foci. Bottom, hierarchical alignment of partitions across the LA and NLA. **f,** Domain graph illustrating spatial adjacency and relationships, separating LA and NLA compartments. **g,** Multi-level spatial ring analysis of Domain 3 foci. Left and middle, global map and enlarged spatial topologies of representative foci (F2 and F7), showing the pure lymphoid core (Inside) surrounded by transition boundaries (Inner and Outer Rings). Right, box plot of local spatial entropy across concentric zones, demonstrating spatial coupling to high entropy remodeling interfaces. **h,** Left, top discriminative gene features learned by stEDGE for Domain 3. Right, spatial co-expression map of the corresponding gene program. **i,** Dot plot of the combined TLS-like gene signature across all domains. Dot size indicates the fraction of expressing cells and color intensity indicates mean expression. **j,** Volcano plot with marginal density distributions comparing differential expression between two spatially distinct lymphoid aggregates (Focus 2 versus Focus 7) within Domain 3, revealing spatial microenvironmental heterogeneity.

To evaluate reconstruction accuracy, we compared stEDGE with seven representative spatial domain methods (Fig. 6c; Supplementary Fig. 80c; Supplementary Table 22). stEDGE demonstrated the highest overall concordance with the reference architecture, ranking first in ARI (0.334), NMI (0.516), and COM (0.470). Although SCAN-IT achieved marginally higher HOM (0.584 versus 0.572), it merged the pleura and expansive interstitium into a single dominant cluster, failing to resolve these as distinct anatomical entities. Consequently, stEDGE achieved the highest overall accuracy score (the mean of ARI, NMI, HOM, and COM). Visually, stEDGE most faithfully preserved the complex multicompartment organization of the fibrotic lung, maintaining the continuity of the broad non-lesion territory alongside interwoven airway, alveolar, and interstitial structures. By contrast, alternative graph- and embedding-based methods suffered from local mixing across these distinct compartments, whereas GASTON produced rigid, axis-like partitions that failed to capture the branching and cavity-rich tissue geometry (Supplementary Fig. 80d). Together, these results show that stEDGE achieves the most balanced reconstruction at single-cell resolution, successfully resolving the trade-off between over-smoothing and artificial fragmentation.

Beyond reconstructing the reference regions, stEDGE organized the Xenium lung section into a nested, multi-level hierarchy spanning fine, domain, and coarse states. While the fine-level resolved 18 spatial microdomains capturing localized remodeling heterogeneity, the domain-level consolidated these states into nine intermediate domains optimized for niche analysis. At the coarsest level, stEDGE partitioned the tissue into three primary structures: a NLA, a lesion-associated fibrotic remodeling area (LA), and a distinct airway-lumen branch (Fig. 6d; Supplementary Fig. 81f–i; Supplementary Fig. 82a,c). Thus, stEDGE represents fibrotic remodeling not as a flat partition, but as a multiscale hierarchy of nested spatial states (Supplementary Fig. 83).

We next leveraged this hierarchy to map domain transition intensity across the lung architecture. The domain-level DTI map revealed a structured transition-stability landscape, showing lower DTI within the NLA and elevated DTI across much of the remodeled LA region (Fig. 6e; Supplementary Fig. 81d). This transition-stability landscape allowed us to identify locally stable spatial states within the remodeling hierarchy (Fig. 6f; Supplementary Fig. 82c). Notably, the airway-lumen branch persisted as an independent coarse compartment across all hierarchical levels, exhibiting exceptionally low DTI relative to other domains (Fig. 6d,e; Supplementary Fig. 81d,f–i; Supplementary Fig. 82a,c).

Tree-guided gene attribution identified *MARCO*^55,55^, *SPP1*, and *AQP9* as key discriminative features of this branch, consistent with a macrophage-associated airway-lumen state^53,56,57^ (Supplementary Fig. 84). These results highlight the ability of stEDGE to preserve highly specific cellular territories rather than forcing localized niches into broader anatomical structures.

Beyond the airway-lumen branch, the transition-stability landscape revealed compact, low DTI foci corresponding to Domain 3 (Fig. 6e). Spatial network analysis confirmed that Domain 3 forms organized, discrete lymphoid aggregates rather than diffuse immune infiltrates, showing a same-domain neighbor fraction of 0.883 (12.3-fold enrichment; Supplementary Fig. 85a,b). Unbiased, tree-guided gene attribution identified a distinct Domain 3 program dominated by *CXCL13*, *MS4A1*, *BANK1*, and *CCL19*^58,59^, indicating a spatially organized, lineage-specific immune state rather than an *a posteriori* marker-defined cluster (Fig. 6h; Supplementary Fig. 86).

To evaluate whether Domain 3 indeed represents a structured TLS-like niche, we performed supervised module scoring using canonical markers for B-cell identity, TLS organization, and a combined TLS-like signature^60–62^ (Supplementary Fig. 87a). Dot plot analysis of the combined signature demonstrated that both the fraction of expressing cells and mean expression were exclusively restricted to Domain 3 (Fig. 6i). Spatial mapping visually confirmed that these active immune programs were tightly localized to the discrete Domain 3 aggregates (Supplementary Fig. 87b). Quantitatively, Domain 3 exhibited prominent, exclusive enrichment of all three scores (Kruskal-Wallis, *P* < 10^−15^; pairwise Mann-Whitney U vs. others: all *P* < 10^−15^, with Cliff’s *δ* = 0.609, 0.561 and 0.646, for B-cell, TLS-organizing, and combined signatures, respectively; Supplementary Fig. 86c). Domain 3 also exhibited near-zero local spatial entropy and significantly reduced PCA-centroid distance compared to other lesion domains (median 2.35 versus 3.33 ; *P* < 0.001), demonstrating high transcriptional coherence and spatial segregation (Supplementary Fig. 85c–f). These spatial and molecular characteristics suggest that Domain 3 likely represents a spatially organized TLS-like lymphoid aggregate niche^58^. Together, these results demonstrate stEDGE’s capacity to resolve complex, pathologically relevant immune microenvironments within highly remodeled tissues. To assess adjacent-section reproducibility, we further analyzed SSc_1_1_1, in which stEDGE independently resolved fine, domain and coarse spatial organization. Fixed TLS-like, B-cell, TLS-organizing and combined TLS signatures were consistently enriched in Domain 4, whose discriminative gene weights showed strong concordance with those of the TLS-like Domain 3 in SSc_1_1_2 (Pearson r = 0.929; Spearman ρ = 0.976; Supplementary Fig. 89a–g).

We next investigated whether these TLS-like foci were isolated immune aggregates or spatially organized relative to the surrounding remodeling interfaces. Graph-based ring analysis around seven robust Domain 3 foci revealed a conserved target-like topology, characterized by a low entropy lymphoid core enveloped by a high entropy inner ring and a gradually transitioning outer ring^63,64^ (Fig. 6g; Supplementary Fig. 88a,b). Local spatial entropy was near zero within the core but increased sharply in the immediate inner ring—a pattern significant at both the single-cell level (Mann–Whitney U test, *P* < 0.001) and the paired-focus level (Wilcoxon signed-rank test, *n* = 7, *P* = 0.0078; Supplementary Fig. 88c,d). Thus, Domain 3 defines stable lymphoid cores embedded within the fibrotic landscape that are spatially coupled to transition-rich tissue interfaces^65^. Consistently, nine TLS-like foci identified in SSc_1_1_1 exhibited TLS-enriched cores surrounded by inner rings with elevated spatial entropy and DTI, supporting adjacent-section reproducibility of the TLS-like core– inner ring organization (Supplementary Fig. 89h,i).

Having established this core–ring organization in the discovery section, we asked whether spatial positioning within the remodeling landscape shapes the transcriptional heterogeneity of these TLS-like niches. We compared Focus 2 (situated at the active LA–NLA boundary) with Focus 7 (embedded deep within the fibrotic LA region; Fig. 6j; Supplementary Data 18). Despite sharing a common Domain 3 identity, the two foci exhibited distinct transcriptional programs tailored to their spatial contexts. Focus 7 was enriched for *CXCL13* together with stromal-network-associated genes (*THY1*, *LTBP2* and *FBN1*), consistent with a lymphoid-recruiting and tissue-organizing program^66^. Conversely, Focus 2 upregulated *MYC*, *PIM1*, *JCHAIN*, *CCL2*, *SELENOP*, and *SERPINA3*, compatible with an interface-exposed activation program with plasma-cell-associated and inflammatory myeloid-crosstalk features^65^. This spatially informed comparison directly links the molecular divergence of TLS-like niches to their microenvironmental coordinates. Collectively, stEDGE resolves fibrotic lung remodeling not as a uniform lesion, but as a coherent, multiscale hierarchy comprising transition-rich lesion interfaces, stable airway-associated macrophage states, and spatially organized yet molecularly heterogeneous TLS-like lymphoid niches.

## Discussion

Spatial transcriptomics has transformed the study of tissue architecture, yet many computational analyses still describe tissues primarily as collections of discrete regions. Our results suggest that this view is incomplete. Across development, inflammation, cancer and brain tissue, stEDGE supports a broader formulation in which spatial organization is jointly defined by stable compartments, transition-rich interfaces, multiscale hierarchy and associated gene programs. In this sense, stEDGE extends spatial analysis beyond domain delineation alone and provides a framework for examining how tissue states are arranged, connected and diversified across space.

A key conceptual advance of stEDGE is that it treats spatial domain boundaries as primary structural signals rather than as implicit consequences of label changes. By integrating boundary probability with domain-level transition propensity and molecular evidence, stEDGE further enables selected boundaries and adjacent states to be interpreted as biological interfaces or transcriptional transition zones. This transition-aware view is integrated with a coarse-to-fine graph and tree organization of spatial states and with tree-guided gene attribution, allowing multiscale structure and multiscale molecular interpretation to be linked within one framework. Together, these components recast spatial reconstruction as a problem of representing not only where domains lie, but also how they transition, branch and acquire distinct molecular identities.

The biological applications of stEDGE point to several broader principles of tissue organization. First, in the analyzed embryo and Crohn’s disease examples, several biologically meaningful interfaces were better represented as directional or finite-width transcriptional transition zones, as illustrated by developmental boundaries in the mouse embryo and ordered epithelial-to-stromal transitions in Crohn’s disease tissue. Second, tissues often comprise stable core compartments together with transition-associated or niche-like local states, rather than uniformly homogeneous regions. Third, higher-order spatial hierarchy can reveal structure that is not reducible to conventional annotation alone, as seen in the hidden neural crest states resolved in the embryo and the hierarchical tumor states identified in breast cancer. These observations argue that compartments, interfaces and local transition states should be regarded as complementary features of tissue architecture rather than alternative descriptions of it.

The Crohn’s disease and breast cancer analyses further illustrate the value of linking multiscale spatial structure to molecular programs. In Crohn’s disease, the combination of ordered transition domains, elevated DTI and localized stromal specialization suggests that inflammatory remodeling is organized not only by broad layered architecture but also by intermediate and niche-like states^67^. In breast cancer, the correspondence between domain hierarchy and node-specific gene programs indicates that tumor organization is more informative when viewed as a structured multiscale system than when reduced to pathology labels alone. More broadly, these findings suggest that biologically consequential states can reside at interfaces, within locally specialized branches or across nested spatial levels that are difficult to recover with conventional partition-based methods. stEDGE therefore provides a common analytical language for studying developmental transitions, inflammatory remodeling and tumor microenvironmental heterogeneity.

The brain analyses extend this perspective by showing that the utility of stEDGE is not confined to one architectural mode. The successful reconstruction of both region-based hippocampal organization and highly folded layer-based cerebellar organization suggests that the framework is structure-agnostic rather than tuned to a single geometric pattern. This is important because ST datasets increasingly span tissues with markedly different organizing principles, from compartmentalized systems to layered, folded or gradient-rich architectures. A useful general framework should therefore accommodate multiple spatial regimes without requiring a separate conceptual representation for each one, and our results suggest that transition-aware multiscale reconstruction provides one such strategy. Several limitations and future directions should be considered. Our current analyses are based primarily on two-dimensional sections, and extension to serial-section, three-dimensional or explicitly temporal spatial datasets may further improve reconstruction of tissue organization. In addition, although tree-guided gene attribution provides an interpretable link between hierarchical spatial structure and molecular programs, further validation using perturbation, lineage-resolved and multimodal spatial datasets will be important for testing the causal basis of these programs. Continued growth in dataset scale and modality diversity will also motivate additional work on computational scalability and integration with morphology-aware and spatial multi-omics frameworks. Despite these limitations, our results support stEDGE as a general strategy for decoding tissue architecture by jointly modeling stable compartments, transition-rich interfaces and multiscale gene programs across complex biological systems.

## Methods

### Data preparation

stEDGE takes as input a spatial transcriptomics dataset consisting of a gene expression matrix *X* and a spatial coordinate matrix *S*. Here, *X* ∈ ℝ*^N^*^×*G*^, where *N* denotes the number of spatial locations and *G* denotes the number of genes, and each entry represents the expression level of a gene at a given location. The matrix *S* ∈ ℝ*^N^*^×2^ records the two-dimensional spatial position of each location. Depending on the experimental platform, a location may correspond to a spot, bin or single-cell. Together, *X* and *S* define the input representation for stEDGE. To ensure general applicability across datasets, all spatial transcriptomics data are converted into this unified matrix-based representation before downstream analysis. No anatomical labels, histological priors or predefined spatial partitions are required as model inputs. When available, reference annotations are reserved for downstream benchmarking and interpretation only.

### Data Preprocessing

Raw expression matrices were filtered, normalized and log-transformed before downstream analysis. Highly variable genes were selected and projected into a low-dimensional space by principal component analysis (PCA), yielding the default transcriptome-based embedding used in stEDGE. For high-resolution datasets, including Slide-seq, MERFISH and cell-level Stereo-seq, stEDGE additionally supports a multiscale neighborhood composition representation. In the default implementation, neighborhood composition is quantified across four spatial scales using an increasing-radius scheme with a base radius of 15, and the resulting features are normalized, log-transformed and projected by PCA to obtain an alternative embedding. This option is designed to better capture local spatial context in near-cellular or single-cell-resolved data.

#### Consensus-based boundary modeling

To explicitly represent tissue interfaces before segmentation, stEDGE first estimates a continuous boundary probability field from multi-resolution consensus structure. Because tissue organization may not be stably captured by a single clustering resolution, stEDGE begins by generating an ensemble of clustering solutions across multiple resolutions or clustering settings. In the default implementation, clustering solutions are generated using the Leiden algorithm on the transcriptome-based embedding over a multi-resolution grid defined by a start value of 0.1, a stop value of 2.0 and a step size of 0.1. stEDGE also supports centroid-based clustering, namely k-means, as an alternative strategy for constructing clustering ensembles under different assumptions. Rather than treating any individual partition as the final structural representation, stEDGE uses recurrent co-assignment patterns across all ensemble solutions to identify local relationships that are stable across scales.

Given a set of *K* clustering solutions 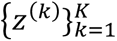, we define the consensus frequency between locations *i* and *j* as the proportion of solutions in which they are assigned to the same cluster:

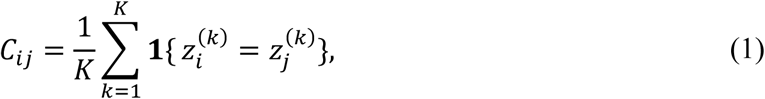

where **1**{⋅} is the indicator function. The diagonal entries of *C* are set to zero in downstream computation. The resulting consensus matrix *C* captures pairwise structural stability across clustering scales, with higher values indicating more stable within-domain relationships and lower values indicating unstable or boundary-adjacent relationships.

To convert this consensus structure into an explicit boundary representation, stEDGE evaluates local consensus consistency within the spatial neighborhood of each location. Let N(*i*) denote the set of spatial neighbors of location *i*. We define the local boundary likelihood as the complement of the average consensus frequency between *i* and its neighbors:

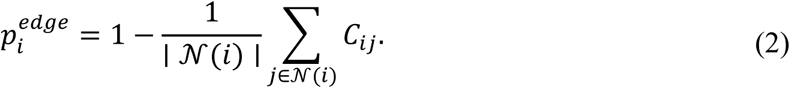

Because *C_ij_* ∈ [0, 1], the raw inconsistency score 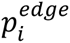 is naturally bounded within [0, 1]. Therefore, stEDGE clips the resulting values to [0, 1] without applying dataset-wise min–max normalization, preserving the absolute magnitude of local boundary uncertainty. Under this formulation, locations with low 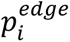 correspond to stable domain interiors, whereas locations with high 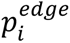 are more likely to reside at tissue interfaces or transition-rich regions. Importantly, this continuous boundary field is not itself a final partition, but instead serves as a structural prior for downstream fine-domain recovery.

### Edge-guided fine-domain discovery

Given the estimated boundary probability field *p^edge^*, stEDGE recovers fine-grained spatial domains through a boundary-preserving expansion procedure. Rather than assigning fine domains directly from a single global clustering resolution, stEDGE first identifies stable local cores in low-boundary regions. In the default implementation, seed candidates are selected as the locations within the lowest 10% of the boundary probability distribution,

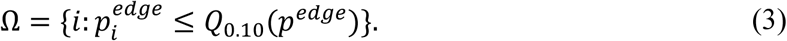

A 6-nearest-neighbor spatial graph is then constructed, and connected components induced by Ωare extracted as initial seed regions S*_r_*. Components with fewer than five locations are discarded, so that the retained seeds represent high-confidence local domain interiors. To guide expansion from these seeds, stEDGE estimates the spatial gradient of the boundary field. Let *x_i_* ∈ ℝ^2^denote the spatial coordinate of location *i*. For each location, the gradient vector *g_i_* is estimated from its spatial neighborhood by weighted local least squares,

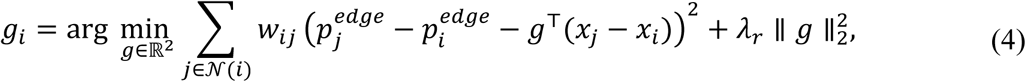

where closer neighbors receive larger weights. In the default implementation, gradients are estimated using 12 spatial neighbors without pre-smoothing of *p^edge^*, preserving narrow boundary signals. The resulting vector field provides a local directional prior that helps avoid expansion toward increasing boundary probability.

Starting from all seed regions, stEDGE performs competitive priority-based region growing. All seeds are expanded simultaneously through a shared priority queue, so neighboring seeds compete for unlabeled locations. Candidate locations are prioritized according to whether the expansion moves toward a boundary, the boundary probability of the candidate, the maximum boundary probability along the local edge, and the local jump in boundary probability. In this way, expansion preferentially proceeds through low-boundary, locally smooth interiors and is delayed or stopped near high-boundary interfaces. A candidate location is accepted only when its boundary probability and local edge-barrier score remain below a seed-specific threshold, and when the local change in *p^edge^*does not exceed a jump threshold. The global boundary threshold is defined as

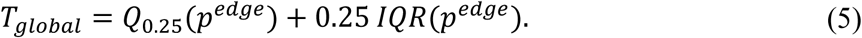

For each seed region, stEDGE also computes a seed-specific threshold from the boundary probabilities within the seed. By default, this seed threshold is capped by *T_global_*, making the expansion conservative and preventing individual seeds from crossing globally strong interfaces. Isolated unlabeled holes surrounded by a single region can be filled if their boundary probability remains below the corresponding threshold. This design allows stEDGE to recover high-confidence domain interiors while remaining conservative near broad or transition-rich tissue interfaces.

Because boundary-preserving growing may initially recover only partial domains, stEDGE further refines intermediate regions using the multi-resolution consensus matrix. For each intermediate region *R_r_*, a consensus support score is computed for every location *j* as

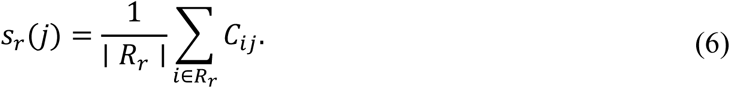

Regions are processed from larger to smaller size, and unassigned locations with *s_r_*(*j*) ≥ 0.7are assigned to the corresponding region. stEDGE further absorbs unassigned locations spatially surrounded by a single region and removes isolated assignments lacking same-region neighbors. Remaining unassigned locations are re-examined by thresholding the consensus matrix within the unassigned subset at 0.7 and extracting connected components with at least 20 locations as additional domains.

Finally, the resulting partial labels are completed by spatially constrained label propagation. stEDGE constructs a joint feature space by concatenating a 50-dimensional PCA representation of the expression matrix with standardized spatial coordinates weighted by 0.3, and then performs label propagation using a 7-nearest-neighbor graph. Original nonzero consensus-region labels are retained, and only unlabeled locations are filled, yielding the final fine-domain assignment 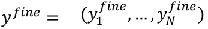.

#### Transition-aware hierarchical refinement of spatial domains

To organize fine domains into higher-order tissue structure, stEDGE performs transition-aware hierarchical refinement based on regional similarity, inter-domain boundaries and transition propensity. Let 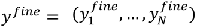 denote the fine-domain assignment obtained in the previous step, and let D = {*d*_1_, …, *d_K_*}denote the set of inferred fine domains. Rather than treating these fine domains as a final flat partition, stEDGE next quantifies how strongly each location is associated with each domain in consensus space, and uses this information to identify both stable compartments and transition-prone states. For each location *i*and domain *d* ∈ D, stEDGE first computes a consensus affinity 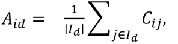, where 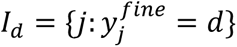 is the index set of locations assigned to domain *d*. These affinities are normalized row-wise to obtain a soft domain assignment matrix

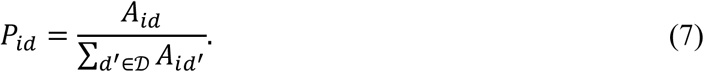

Using this soft assignment, stEDGE defines the spot-level transition score as the normalized Shannon entropy

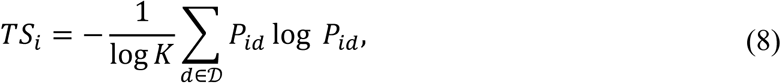

with *TS_i_* = 0 when *K* = 1. Under this formulation, *TS_i_* approaches 0 when location *i* is strongly associated with a single domain, and increases toward 1 when its consensus affinities are distributed across multiple domains. The domain transition index is then defined as the average spot-level transition score within each domain:

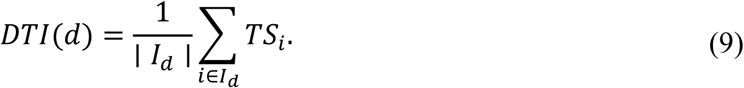

Thus, domains with low DTI correspond to relatively stable compartments, whereas domains with high DTI are more transition-prone or structurally mixed. The same soft assignment matrix is used to define inter-domain similarity: 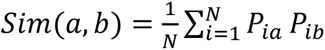. In parallel, stEDGE constructs a 6-nearest-neighbor spatial graph over all locations and records only cross-domain contacts to define domain adjacency. For each adjacent pair (*a*, *b*), boundary strength is quantified as the mean boundary probability across contacting cross-domain neighbor pairs, Bound 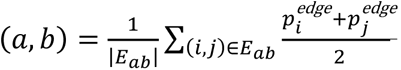, where *E_ab_* denotes the set of spatial neighbor pairs with 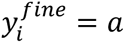 and 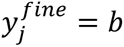 This yields three complementary quantities for hierarchical refinement: domain similarity, transition propensity and inter-domain boundary strength.

stEDGE then performs iterative domain merging under a transition-aware criterion. Starting from singleton fine-domain clusters, each candidate merge is evaluated by aggregating inter-cluster similarity, cluster-level DTI and boundary strength across the fine domains contained in the two clusters. Inter-cluster similarity is computed as a size-weighted average of pairwise domain similarities, cluster-level DTI is computed as a size-weighted average of domain-level DTI values, and inter-cluster boundary strength is computed from the boundary strengths of adjacent domain pairs.

To make the merge score comparable across datasets, stEDGE standardizes the similarity, stability and boundary terms using the distribution of initial fine-domain pairs. The merge score is defined as

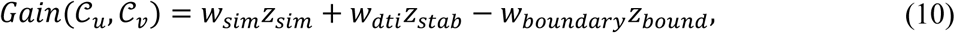

where *z_sim_*, *z_stab_* and *z_bound_* are the standardized inter-cluster similarity, stability and boundary-strength terms, respectively. Stability is defined from DTI so that lower transition propensity gives a higher stability contribution. By default, *w_sim_* = 1.0, *w_dti_* = 0.5, *w_boundary_* = 0.5, and the minimum gain threshold is 1.0. A conservative post-merge quality check further prevents merging when the predicted merged DTI becomes too high.

At each iteration, stEDGE merges the candidate pair with the highest gain until no candidate exceeds the minimum gain threshold or a target number of domains is reached. The merged partition defines the intermediate domain level, denoted by 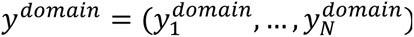. After domain merging, stEDGE recomputes transition scores and domain similarity for the merged domain level. It then constructs a domain-level weighted graph from the post-merge similarity matrix. Weak edges are removed using an adaptive similarity quantile threshold rather than a fixed similarity cutoff; by default, edges below the median off-diagonal similarity are pruned. Leiden clustering is then applied to this domain-level graph at resolution 1.0 to obtain coarse domains, 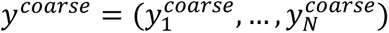Together, *y^fine^*, *y^domain^*and *y^coarse^*define a three-level hierarchy of multiscale tissue structure. **Tree-guided multiscale gene attribution.** To interpret molecular programs in the same multiscale context in which spatial states are organized, stEDGE introduces a tree-guided gene attribution framework. A central challenge in multiscale spatial analysis is that genes defining a broad parent compartment are often distinct from those that resolve divergence among its child branches. Consequently, molecular interpretation cannot be reduced to marker identification on flat partitions alone, because such analyses tend to conflate shared parent-level identity with branch-specific specialization. To address this, stEDGE performs gene attribution directly on the reconstructed hierarchy.

Using the three-level assignments *y*^fine^, *y*^domain^and *y*^coarse^, stEDGE constructs a rooted hierarchy T = {root → coarse → domain → fine}. For each internal node *v* ∈ T, let Ch(*v*) denote its child nodes and ℐ*_v_* the set of locations descending from *v*. Each child *u* ∈ Ch(*v*) defines a local node-split task on ℐ*_v_*, in which locations belonging to *u*are treated as positives and those assigned to the remaining children of *v* are treated as negatives:

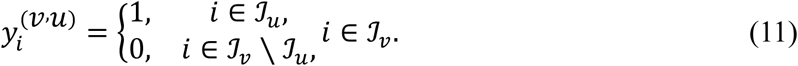

Let *T* denote the total number of node-split tasks and *W* ∈ ℝ*^T^*^×*G*^ the corresponding task-by-gene weight matrix, where the *t*-th row *w_t_* encodes the gene weights for task *t*. stEDGE estimates *W* by jointly optimizing a fitting term together with tree-guided and sibling-exclusive regularization:

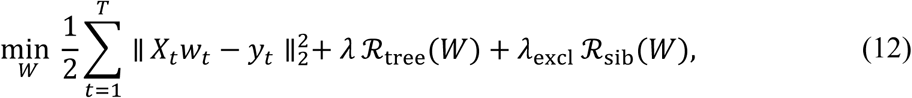

where *X_t_* is the expression matrix restricted to the samples involved in task *t*, and *y_t_* is the corresponding binary label vector. The tree-guided term encourages tasks within the same subtree to share genes,

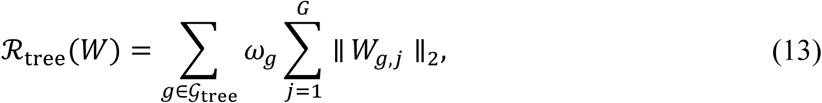

where G_tree_ denotes the collection of subtree task groups, *W_g_*_,*j*_ is the subvector of weights for gene *j*across the tasks in group *g*, and *ω_g_* is a group weight. This term reflects the idea that descendant spatial states often retain a shared transcriptional backbone inherited from their parent compartment. To separate this shared identity from local branch divergence, stEDGE further imposes a sibling-exclusive penalty,

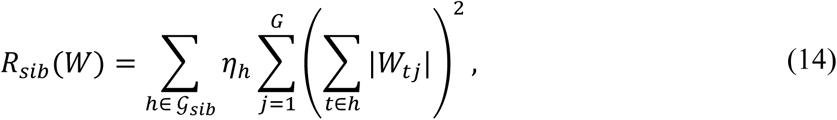

where G_sib_ denotes the collection of sibling task groups branching from the same parent node, and *η*_ℎ_ is a group-specific weight. This term discourages repeated use of the same genes across sibling branches, because genes uniformly shared across all children are more appropriately interpreted as parent-level programs rather than branch-specific determinants. In the default implementation, stEDGE uses the top 3,000 highly variable genes and optimizes the model with *λ* = 0.2, *λ*_excl_ = 0.1, 200 iterations and a convergence tolerance of 10^−4^.

After optimization, multiscale gene programs are read out from *W*. A global gene score is defined

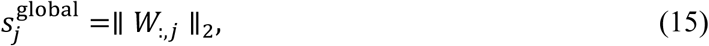

which summarizes the overall contribution of gene *j* across all node-split tasks. Node-level programs are obtained by aggregating weights across tasks descending from the same internal node. Branch-level programs are obtained from individual node-split tasks and ranked by absolute task-specific gene weights, while the signed weights indicate whether a gene contributes positively or negatively to a branch distinction. These two readouts respectively capture shared parent-level identity and child-specific divergence. By preserving this distinction, stEDGE interprets molecular programs in a hierarchy-aware rather than flat-cluster framework.

### Usage of Comparative Methods

To comprehensively evaluate the performance of stEDGE, we compared it with representative spatial transcriptomics methods, including **stLearn** (https://github.com/BiomedicalMachineLearning/stLearn), **SEDR** (https://github.com/JinmiaoChenLab/SEDR/), **STAGATE** (https://github.com/zhanglabtools/STAGATE), **STAIG** (https://github.com/y-itao/STAIG), **GraphST** (https://github.com/JinmiaoChenLab/GraphST), **PROST**(https://github.com/Tang-Lab-super/PROST), **SCAN-IT**(https://github.com/zcang/SCAN-IT), **SpaGCN** (https://github.com/jianhuupenn/SpaGCN), **STAMP** (implemented via scTM: https://github.com/JinmiaoChenLab/scTM), **GASTON** (https://github.com/raphael-group/GASTON), and **SAGE** (https://github.com/yihe-csu/SAGE). In comparative experiments, for datasets with available reference annotations, the number of inferred clusters for each method was set to match the number of annotated domains to ensure a fair comparison across methods. Unless otherwise specified, all methods were run using their recommended or default parameter settings and preprocessing pipelines as described in their original publications or official implementations. To ensure reproducibility, a fixed random seed was used across all experiments.

### Performance Evaluation Metrics

We evaluated spatial domain reconstruction and boundary localization using complementary clustering, spatial continuity and interface-specific metrics. Domain agreement with reference annotations was quantified using adjusted rand index (ARI), normalized mutual information (NMI), homogeneity (HOM) and completeness (COM), where higher values indicate better agreement. For datasets without reference annotations (for example, mouse hippocampus and cerebellum), clustering quality was additionally assessed using silhouette score and Davies-Bouldin index (lower is better), together with spatial variants including spatial silhouette and spatial coherence to quantify joint transcriptomic–spatial consistency. Spatial continuity was assessed using ASW, CHAOS and PAS; ASW evaluates spatial separation between domains (higher is better), whereas CHAOS and PAS quantify local spatial inconsistency (lower is better). Boundary localization was evaluated by comparing predicted boundary scores with ground-truth interfaces derived from reference annotations. Spatial neighbor pairs with different reference labels were defined as boundary edges. Node-level boundary scores were projected to edges by averaging the scores of connected locations. Performance was quantified using AUPRC and AUROC (higher is better), as well as F1@q95, precision (P@q95) and recall (R@q95), where *q*_95_ denotes the 95th percentile of predicted boundary scores. To characterize transcriptional transitions, we performed signed distance profiling relative to domain interfaces and quantified transition breadth as the spatial extent of continuous changes across boundaries.

#### Boundary evaluation and interface analysis

We use “domain boundary” for the annotation-derived or model-inferred separation between adjacent domains. Selected boundaries are referred to as biological interfaces only when they correspond to a defined anatomical or pathological juxtaposition, and as transcriptional transition zones when continuous distance-dependent molecular variation is observed. For datasets with available reference annotations, ground-truth boundary edges were defined on the spatial neighbor graph. Specifically, for each pair of spatially adjacent locations (*i*, *j*), an edge was labeled as a boundary edge if the reference labels differed 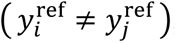, and as a within-region edge otherwise 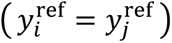. To compare node-level boundary predictions with these edge-level labels, boundary probabilities 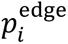 were projected to edges by averaging the scores of connected locations, 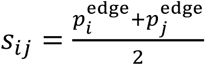 These edge scores were then used to quantify boundary localization performance as a binary classification problem between boundary and within-region edges. To further characterize interface structure, we analyzed transcriptional transitions across selected domain boundaries using signed distance profiling. For each interface, spatial locations were assigned a signed distance relative to the boundary, and gene expression or domain assignment was examined as a function of this distance. Transition breadth was defined as the spatial extent over which these quantities varied continuously across the interface, providing a measure of whether boundaries correspond to sharp separations or gradual transitions.

### Identification of differential genes

Differentially expressed genes (DEGs) were identified using a two-sided Wilcoxon rank-sum test as implemented in the SCANPY Python package (version 1.10.2). Unless otherwise specified, comparisons were performed between groups of spatial locations defined by domain, subdomain or interface assignments. *P* values were adjusted for multiple-testing using the Benjamini–Hochberg procedure, and genes were considered significant at a false discovery rate (FDR) threshold of 0.005. In addition, we required an absolute log_2_ fold change of at least 1 (∣ log _2_ FC ∣≥ 1) between groups. Fold changes were computed from the mean expression of each gene within each group. In addition to pairwise differential expression analysis, stEDGE further enables identification of node-specific gene programs in the reconstructed hierarchy. For each internal node, genes are ranked according to their ability to distinguish one branch from its siblings within the local context of descendant spatial locations. By conditioning comparisons on the hierarchical structure, this formulation separates branch-specific specialization from globally shared expression patterns and yields gene programs associated with multiple levels of spatial organization. Unlike conventional differential expression analysis, this hierarchy-aware attribution does not rely on explicit statistical hypothesis testing; instead, genes are prioritized based on their contribution to discriminating spatial states within the inferred hierarchy.

### Gene Ontology functional enrichment analysis

Gene Ontology (GO) and pathway enrichment analyses were performed using the *enrichr* function from the *gseapy* Python package (version 1.1.4). For mouse embryo datasets (Data 1–9), enrichment was conducted using the KEGG_2019_Mouse and GO_Biological_Process_2025 libraries. For Crohn’s disease datasets (Data 10–11), GO_Biological_Process_2025 was used. For the human breast cancer dataset (Data 12), KEGG_2021_Human and GO_Biological_Process_2025 were applied. For mouse brain datasets (Data 13–14), enrichment was performed using KEGG_2019_Mouse, GO_Biological_Process_2023 and WikiPathways_2019_Mouse. Enrichment results, including terms, adjusted P values and associated gene sets, are provided in Supplementary Data 6–7 (mouse embryo), Supplementary Data 11 (Crohn’s disease), Supplementary Data 13 (human breast cancer) and Supplementary Data 15 and 17 (mouse brain).

#### Manual reference annotation of the Xenium lung section

We analyzed SSc_1_1_2, a Xenium section from the publicly available SSc-ILD lung dataset generated by Markov et al. Benchmark annotations were manually curated at structural and pathological levels. Structural annotation was performed in QuPath v0.7.0 by integrating nuclear and multiplexed immunofluorescence images with tissue morphology to delineate major anatomical compartments, including airway, alveolar, vascular, interstitial and pleural regions. Because small vascular profiles, alveolar structures and locally remodeled interstitial regions were difficult to define by morphology alone, we further used transferred cell-type labels and known marker-gene patterns as auxiliary evidence for boundary refinement. For cell-type-assisted refinement, we used a published human lung single-cell reference from Natri *et al*.^68^ Fine-grained reference annotations were collapsed into broad lung cell classes, including epithelial, stromal, endothelial/vascular, myeloid and lymphoid lineages. Broad cell-type labels were transferred to Xenium cells using Seurat anchor-based label transfer with SCT normalization. These labels were used only to guide manual refinement of anatomical structures and were not used as inputs to stEDGE. A coarse pathological annotation was generated by manually separating the section into non-lesion-like area (NLA) and lesion-associated fibrotic remodeling area (LA), based on tissue morphology, cell density, cell-type composition and fibrotic remodeling. The resulting annotations were used only as benchmark references.

### Ablation study

To quantitatively evaluate the contribution of individual stEDGE modules, we tested six ablation variants across nine independent spatial transcriptomics datasets (Supplementary Fig. 90, Supplementary Fig. 91 and Supplementary Table 23). The complete baseline model achieved a mean ARI of 0.3719. The results show that removing the spatial graph-driven edge expansion (ABL2) had the most severe impact on model performance, causing a 51.1% decrease in the mean ARI (ΔARI = −0.190). Removing the multi-resolution consensus mechanism (ABL1) resulted in the second-largest decline (ΔARI = −0.069). Eliminating the hierarchical merging constraint (ABL5) improved Homogeneity (ΔHOM = +0.046). However, it significantly increased the predicted number of domains (*N*_pred_), leading to spatial over-fragmentation and a subsequent reduction in both Completeness (ΔCOM = −0.024) and overall ARI. Additionally, removing the spatial and boundary weights (ABL6) also decreased clustering quality (ΔARI = −0.038). In the evaluation of the tree-guided multiscale gene attribution module (Supplementary Fig. 92 and Supplementary Table 24), although the overall Edge AUC showed no significant difference across variants (e.g., remaining constant at 0.8904 in Data_1), omitting the exclusive lasso penalty (ABL8) increased the gene overlap between sibling branches (Sibling Jaccard similarity) by up to 4.4%.

### Parameter robustness

We conducted a sensitivity analysis on the core parameters of stEDGE across the nine datasets (Supplementary Fig. 93 and Supplementary Tables 27–30). For the spatial layout parameters, varying the seed quantile (seed_quantile, 0.05–0.30) and the global threshold *k* (global_thr_k, 0.0–1.0) resulted in minimal fluctuations in the ARI, demonstrating low sensitivity to initialization parameters. For the cluster merging parameters, model performance peaked within specific intervals for the consensus threshold (0.5–0.7) and minimum modularity gain (min_gain, 0.5– 2.0). Under the default parameter settings (consensus threshold = 0.7, min_gain = 1.0), the mean predicted domain count (*N*_pred_) curves across datasets intersected the ground-truth baseline (*N*_gt_ ≈ 14). Sparsity parameter testing within the tree-guided multiscale gene attribution module (Supplementary Fig. 92 and Supplementary Tables 25–26) revealed that increasing lam from 0.01 to 0.50 reduced the number of retained non-zero genes by 49.2% (from 3,000 to 1,524), while the Edge AUC was maintained at 0.8718. Furthermore, increasing the exclusive lasso parameter (*lam_excl*) led to a monotonic 12.2% decrease in sibling Jaccard similarity, with the Edge AUC remaining constant.

#### Statistical analysis Statistics & Reproducibility

All statistical tests and multiple-testing corrections follow the procedures described in the corresponding Methods subsections. In particular, DEG are defined as described in “Identification of differential genes,” with FDR-controlled Wilcoxon rank-sum tests and consistent fold change thresholds across analyses. No statistical method is used to pre-determine sample size. No data is excluded from the analysis, all genes in datasets are used throughout all analyses. The investigators are blinded to allocation during experiments and outcome assessment.

### Runtime and memory profiling

Runtime profiling was performed on a dual-socket Intel Xeon Gold 6230R CPU workstation with 52 physical cores and 104 logical threads, without GPU acceleration. For the main profiling analysis, stEDGE was run on 17 ST datasets using 32 CPU threads, with three independent runs per dataset. Full stEDGE runtime was defined as boundary modeling, fine-domain reconstruction, hierarchical refinement and tree-guided gene attribution, excluding file loading and visualization. Across datasets ranging from 2,541 to 82,224 cells or spots, full runtime ranged from 29.8 to 353.4 s, with a median of 102.5 s; peak RAM ranged from 1.44 to 54.15 GB (Supplementary Fig. 94 and Supplementary Table 31). Runtime and memory scaled strongly with dataset size, and boundary modeling was the dominant runtime component in most datasets. Thread-scaling experiments across 8, 16, 32 and 64 CPU threads showed that 64 threads achieved the shortest runtime in 15 of 17 datasets, with a median 1.38-fold speedup relative to 8 threads, while peak RAM remained similar across thread settings (Supplementary Fig. 95 and Supplementary Table 32).

## Data availability

Publicly available spatial transcriptomics datasets used in this study are summarized in Supplementary Tables 1–2. Mouse embryo datasets (Data 1–9) were obtained from the MOSTA atlas (Chen et al., Cell 2022) and are available at the China National GeneBank Database (CNGBdb) via https://db.cngb.org/stomics/mosta/. Human intestinal datasets from Crohn’s disease patients (Data 10– 11) were obtained from Kong et al. (Nature Genetics 2025). Human breast cancer datasets (Data 12 and Data 16) were obtained from 10x Genomics public resources, including two 10x Visium sections from a publicly available human breast cancer dataset. Mouse brain datasets (Data 13–14; hippocampus and cerebellum) were obtained from Slide-seqV2 (Stickels et al., Nature Biotechnology 2021) and are available via the Broad Institute Single-Cell Portal at https://singlecell.broadinstitute.org/single_cell/study/SCP815. Xenium datasets of systemic sclerosis-associated interstitial lung disease (SSc-ILD) (Data 15 and Data 17; human fibrotic lung sections SSc_1_1_2 and SSc_1_1_1) were obtained from Markov et al.^53^ and are available from NCBI GEO under accession GSE303048.

## Code availability

stEDGE is publicly available at GitHub [https://github.com/yihe-csu/stEDGE].

## Funding

This work was supported in part by the National Key Research and Development Program of China (No.2021YFF1201200), the National Natural Science Foundation of China under Grants (Nos. 62350004, 62332020). This work was carried out in part using computing resources at the High-Performance Computing Center of Central South University.

## Author contributions

Y.H. and S.W. conceived the study. Y.H. designed the stEDGE framework, developed the computational method, implemented the software, performed the spatial transcriptomics analyses, benchmarking experiments and ablation studies, generated the figures and wrote the initial manuscript draft. S.C. contributed to data collection, preprocessing, comparative analyses and result interpretation. L.D., R.H. and H.Z. contributed to sample annotation, biological interpretation and result validation. X.P. and G.D. contributed to data organization, analysis checking and manuscript revision. H.-D.L. and J.W. contributed to methodological discussion, computational analysis design and interpretation of the results. S.W. and J.W. supervised the project, provided overall guidance on method development and manuscript preparation, and acquired funding. All authors discussed the results, reviewed and edited the manuscript, and approved the final version.

## Competing interests

The authors declare no competing interests.

## Notes

### Competing Interest Statement

The authors have declared no competing interest.

